# Identification and characterization of distinct murine brown adipocyte lineages

**DOI:** 10.1101/2020.08.24.264416

**Authors:** Ruth Karlina, Dominik Lutter, Viktorian Miok, David Fischer, Irem Altun, Theresa Schöttl, Kenji Schorpp, Andreas Israel, Cheryl Cero, James W. Johnson, Ingrid Kapser-Fischer, Anika Böttcher, Susanne Keipert, Annette Feuchtinger, Elisabeth Graf, Tim Strom, Axel Walch, Heiko Lickert, Thomas Walzthoeni, Matthias Heinig, Fabian J. Theis, Cristina García-Cáceres, Aaron M. Cypess, Siegfried Ussar

## Abstract

Brown adipose tissue (BAT) plays an important role in the regulation of body weight and glucose homeostasis. While increasing evidence supports white adipose tissue heterogeneity, little is known about heterogeneity within murine BAT. Using single cell RNA sequencing of the stromal vascular fraction of murine BAT and analysis of 67 brown preadipocyte and adipocyte clones we unravel heterogeneity within brown preadipocytes. Statistical analysis of gene expression profiles from these clones identifies markers distinguishing brown adipocyte lineages. We confirm the presence of distinct brown adipocyte populations *in vivo* using three identified markers; Eif5, Tcf25, and Bin1. Functionally, we demonstrate that loss of Bin1 enhances UCP1 expression and mitochondrial respiration, suggesting that Bin1 marks a dormant brown adipocyte type. The existence of multiple brown adipocyte lineages suggests distinct functional properties of BAT depending on its cellular composition, with potentially distinct function in thermogenesis and the regulation of whole body energy homeostasis.

## Introduction

Brown adipose tissue (BAT) is a comparably small adipose tissue depot. Yet, in light of pandemics of obesity and the metabolic syndrome, much attention is focused on the ability of BAT to dissipate energy in form of heat through uncoupling protein-1 (Ucp1) mediated mitochondrial uncoupling (*1, 2*). Increasing BAT activity results in weight loss promoting metabolic health (*3, 4*), whereas reduction in BAT mass or function associates with increased adiposity and the metabolic syndrome (*5, 6*). In mice, classical BAT depots are found predominantly in the interscapular region as well as around the neck (*7*). Murine BAT development starts around embryonic days 9.5-11.5 (*8*), through differentiation of a Myf5 positive skeletal muscle/ brown adipocyte precursor population (*9*). In contrast, most murine white adipose tissues (WAT) appear largely after birth. Myf5 positive precursor cells were long considered to only differentiate into murine brown adipocytes. Recently, however, expression of Myf5 was also found in white adipocyte precursor cells of different WAT depots (*10-12*), questioning the selectivity of Myf5 positive precursors for the brown adipocyte lineage. Sebo and colleagues, reported that in addition to Pax3^+^/Myf5^+^ cells, a minor fraction of interscapular brown adipocytes derive from the central dermomyotome expressing Pax7 and Myf5 (*13*). Thus, these data suggest that more than one brown adipocyte precursor cell population might exist, albeit the current dogma suggests a single precursor population (*12*).

In addition to classical BAT, chronic cold exposure, β-adrenergic agonist treatment, as well as other factors, such as nutrients, trigger the formation of so called beige, brown-like, adipocytes in murine WAT (*14, 15*). These cells are similar to brown adipocytes, expressing Ucp1 and dissipating energy through mitochondrial uncoupling, but differ in gene expression (*16, 17*) and specific functions (*18, 19*) from classical brown adipocytes. Detailed understanding of the developmental origins of murine brown and beige adipocytes is essential to foster the translation of basic murine research data to human physiology, as the nature of adult human brown adipocytes remains a matter of active debate. Comparative gene expression analysis initially suggested that human brown adipocytes resemble a gene expression signature more closely related to murine beige adipocytes (*17, 20, 21*). However, additional studies did not observe this correlation and suggested higher similarities between murine and human brown adipocytes (*16*). Moreover, several human BAT depots exist that show different histological and functional features and potentially distinct developmental origins (*22-24*). Indeed, Xue and colleagues described heterogeneity in Ucp1 expression in immortalized single cell clones derived from human BAT (*25*). However, it remains to be determined if this heterogeneity depends on genetic differences between donors, environmental factors or the existence of distinct human brown adipocyte lineages. The latter is appealing, as the existence of multiple human brown adipocyte lineages, and potential differences in the inter-individual contribution of these lineages to human BAT, could explain some of the discrepancies in the comparison of human BAT to murine beige and brown adipocytes. These findings also raise the question if murine BAT is, as commonly believed, composed of one brown adipocyte lineage, or a mixture of brown adipocytes derived from multiple lineages. Indeed, housing mice at thermoneutrality, mimicking a physiological state more closely related to humans, results in murine brown adipose tissue resembling many of the histological and functional features of human BAT (*26*). Importantly, murine BAT at thermoneutrality also presents histologically as a mixture of unilocular and multilocular adipocytes, resembling white and brown/ beige adipocytes, respectively. The heterogeneous response of murine BAT in response to changes in ambient temperature further supports functional or developmental heterogeneity among brown adipocytes within one depot. This was recently supported by Song and colleagues, who identified using genetic lineage tracing, scRNAseq and other methods, the presence of Ucp1 low and high murine brown adipocyte subtypes within interscapular brown adipose tissue (*27*).

Similar to this, we observe heterogeneity in Ucp1 expression of murine interscapular BAT *in vivo*. Using single cell RNA sequencing (scRNAseq), we characterize the composition of the stromal vascular fraction of murine BAT. Computational analysis of the scRNAseq data, however, revealed that identified brown preadipocyte clusters represent different stages of adipocyte differentiation rather than distinct preadipocyte lineages. Using an alternative approach to identify distinct brown adipocyte lineages in mice, we performed a detailed phenotypic and computational analysis of brown preadipocyte and adipocyte clones derived from the interscapular murine BAT. Using nonlinear manifold learning approaches we identify a set of potential brown adipocyte lineage markers with distinct correlations to various known markers of BAT. We identify Eif5, Tcf25, and Bin1 as markers for brown adipocyte subsets, as well as their precursors, each one labeling between ∼25-35% of brown preadipocytes and adipocytes *in vivo*. Functionally we demonstrate that loss of Bin1 increases thermogenic gene expression and mitochondrial respiration. Thus, our data strongly support the existence of multiple murine brown adipocyte lineages and identify Bin1 as a negative regulator of thermogenesis in brown adipocytes.

## Results

### scRNAseq reveals composition and heterogeneity within the stromal vascular fraction of murine BAT

Ucp1 is the most specific and selective marker for murine and human BAT (*28*). Comparison between interscapular and subscapular BAT in adult male C57Bl/6 mice, did not show differences in Ucp1 expression between these depots (**Supplementary Fig. 1A**). However, as recently reported (*27*), immunofluorescence stainings of adult murine interscapular BAT revealed a mosaic of Ucp1 expression throughout the tissue (**Fig. 1A**). Quantification of Ucp1 immunoreactivity and grouping into quartiles of Ucp1 content showed an even split between high and low Ucp1 expressing brown adipocytes (**Fig. 1B, Supplementary Fig. 1B**). To obtain insights into composition and heterogeneity of the stromal vascular fraction (SVF) of BAT, including brown preadipocytes, we performed scRNAseq from the SVF of interscapular BAT of 8 week old male C57BL/6J mice. Louvain clustering of the gene expression of 4013 cells identified 12 distinct clusters (**Fig. 1C** and **Supplementary Data 1**). Using sets of marker genes we identified several immune cell populations (**Fig. 1C** and **Supplementary Data 1**). Moreover, using Pdgfra expression, as well established marker for preadipocytes, we identified two distinct brown preadipocyte clusters (**Fig. 1C** and **Supplementary Fig. 1C** and **Supplementary Data 1**). We also identified a putative mature brown adipocyte cluster (cluster 7) with high Fabp4 and Pparg expression (**Fig. 1C** and **Supplementary Fig. 1C**), which is further supported by the high count of mitochondrial genes in this cluster (**Supplementary Data 1**). Analysis the preadipocytes only (clusters 0 and 8) identified four clusters (**Fig 1D**). Differential gene expression and KEGG enrichment analysis identified a number of differentially expressed genes and related pathways (**Fig. 1E** and **Supplementary Table 1**). Clusters 0,1 and 2 were assigned with a number of distinct pathways, whereas cluster 3 shared most of its enriched pathways except ‘Wnt signaling’ with other clusters. Cluster 0 associated with PPAR and Apelin signaling, both important for terminal brown adipocyte differentiation (*29, 30*), whereas clusters 1-3 showed an enrichment in pathways, such as Wnt, TNF, relaxin and TGFb signaling that are generally associated with the inhibition of adipogenesis (*31-33*). Interestingly, cluster 3 shared most pathways with other clusters, regardless of their distance in the louvain plots. Pseudotime mapping of the preadipocytes largely replicated the louvain clustering, albeit it revealed two distinct developmental branches, with one common precursor (**Fig. 1F**). Analysis of the expression of known markers during adipogenesis identified cluster 2 (enriched in Cd34 expressing cells) as the most undifferentiated state, with transition states (expressing Cebpd and Cebpb) representing clusters 1 and 3. Cluster 0, as suggested by the KEGG enrichment analysis, showed enrichment in cell expressing terminal differentiation markers such as Cebpa, Pparg, Fabp4 and Lpl (**Fig. 1G**). Thus, these data strongly suggest that the four clusters identified by scRNAseq do not represent individual preadipocyte lineages, but rather distinct differentiation stages of preadipocytes. This further suggests that the differences in gene expression upon differentiation mask differences in gene expression between potential brown preadipocyte lineages.

**Figure 1:**
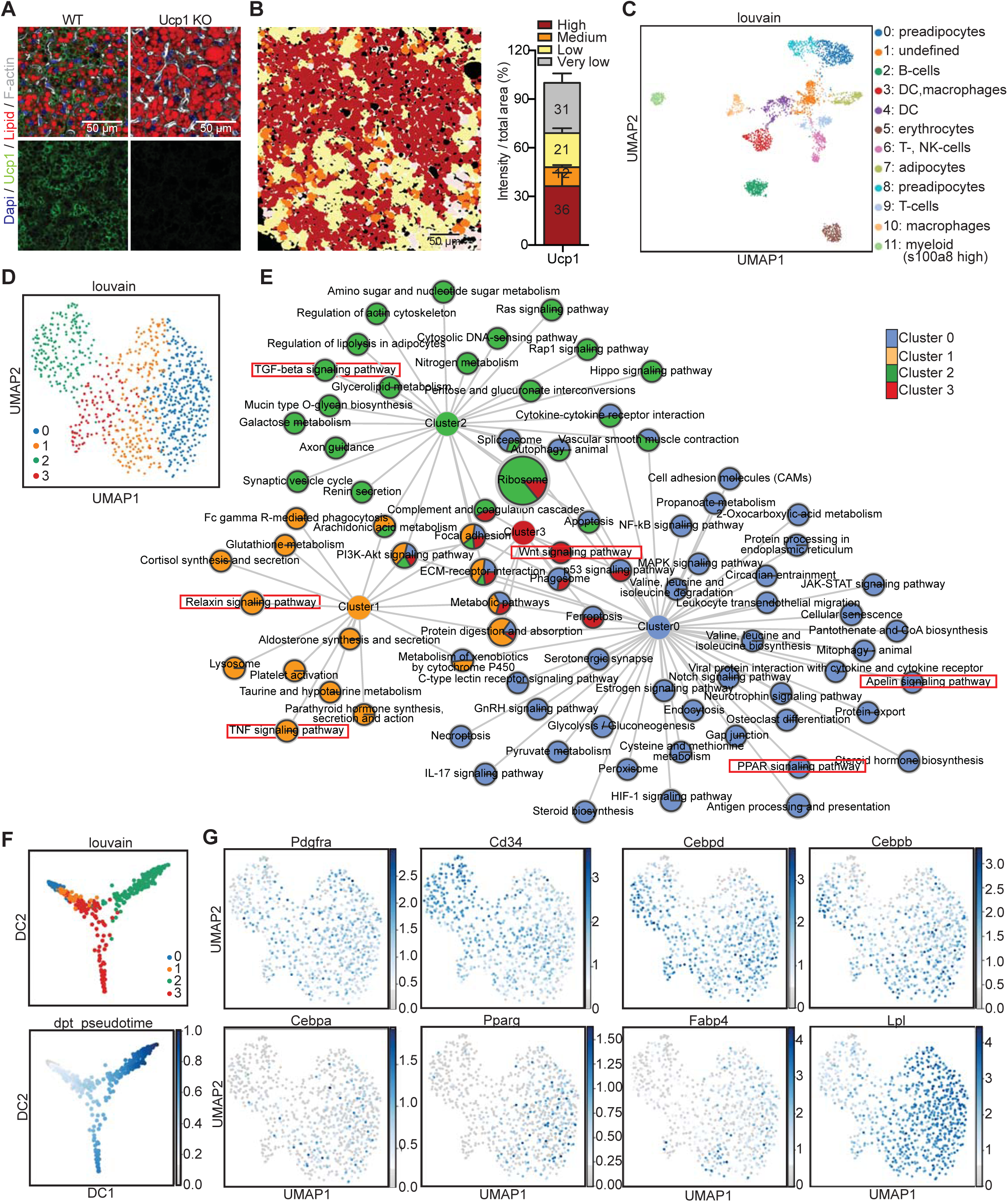
scRNAseq identifies distinct stages of brown adipocyte differentiation. (**A**) Immunofluorescence staining of Ucp1 (green), F-actin (white), lipid droplets (red) and DAPI (blue) from WT and Ucp1 knock-out mice. (**B**) Quantifications of Ucp1 content in individual brown adipocytes in BAT normalized to total area (right panel, n= 11). Lipid droplets are included into area measured and F-actin was used to distinguish each cell. (**C**) UMAP computed on full processed single-cell RNA-seq data set with annotated louvain clusters superimposed. (**D**) UMAP computed on set of preadipocytes with louvain sub-clustering. (**E**) Network of KEGG enriched pathways of preadipocyte clusters. Significantly enriched pathways are connected to the respective cluster nodes. Size of the Pathway nodes and proportion of the node-pie-chart refer to –log10 of enrichment p-value. Node colors refers to single cell cluster identified in (D). (**F**) Diffusion map dimensionality reduction of preadipocytes colored by Louvain clusters (top panel) and pseudotime (lower panel). (**G**) Expression of Pdgfra, Cd34, Cebpd, Cebpb, Cebpa, Pparg, Fabp4 and Lpl in preadipocytes shown in (D). The expression values are size factor normalized and log transformed.

### Murine brown adipocyte clones differ in brown, beige and white marker gene expression

To overcome the limitations of a static experimental setup such as scRNAseq, we established 67 clonal cell lines, derived from the SV40 large-T immortalized stromal vascular fraction of three eight-week old male C57Bl/6 mice. Most cell lines, when differentiated for eight days, accumulated lipids, albeit to different extents, as quantified by Oil Red O (ORO; **Fig. 2A**) and LipidTOX stainings (**Fig 2B** and **C**). Morphological analysis of the preadipocyte and adipocyte clones using LipidTOX and phalloidin (to stain the F-actin cytoskeleton) did not reveal major differences in shape or lipid droplet size between the clones, with the exception of clones 1A6 and 1D5 which had smaller lipid droplets when correlated with the overall lipid content (**Fig. 2C**). Correlation between lipid content (ORO) and Pparg expression at day 8 of differentiation (**Fig. 2D**) showed a strong correlation between lipid accumulation and Pparg expression. However, there were several clones (1B1, 1B3 and 1B6, 2C12, 3C7 and 3D6) that diverged and showed higher Pparg expression than anticipated from the lipid content. In line with the function of Pparg in driving brown adipocyte selective gene expression, clones 1B3, 1B6, 2B11, 2C7, 2C12, 3C7 and 3D6 showed highest Ucp1 expression compared to all other clones (**Fig. 3A**).

**Figure 2:**
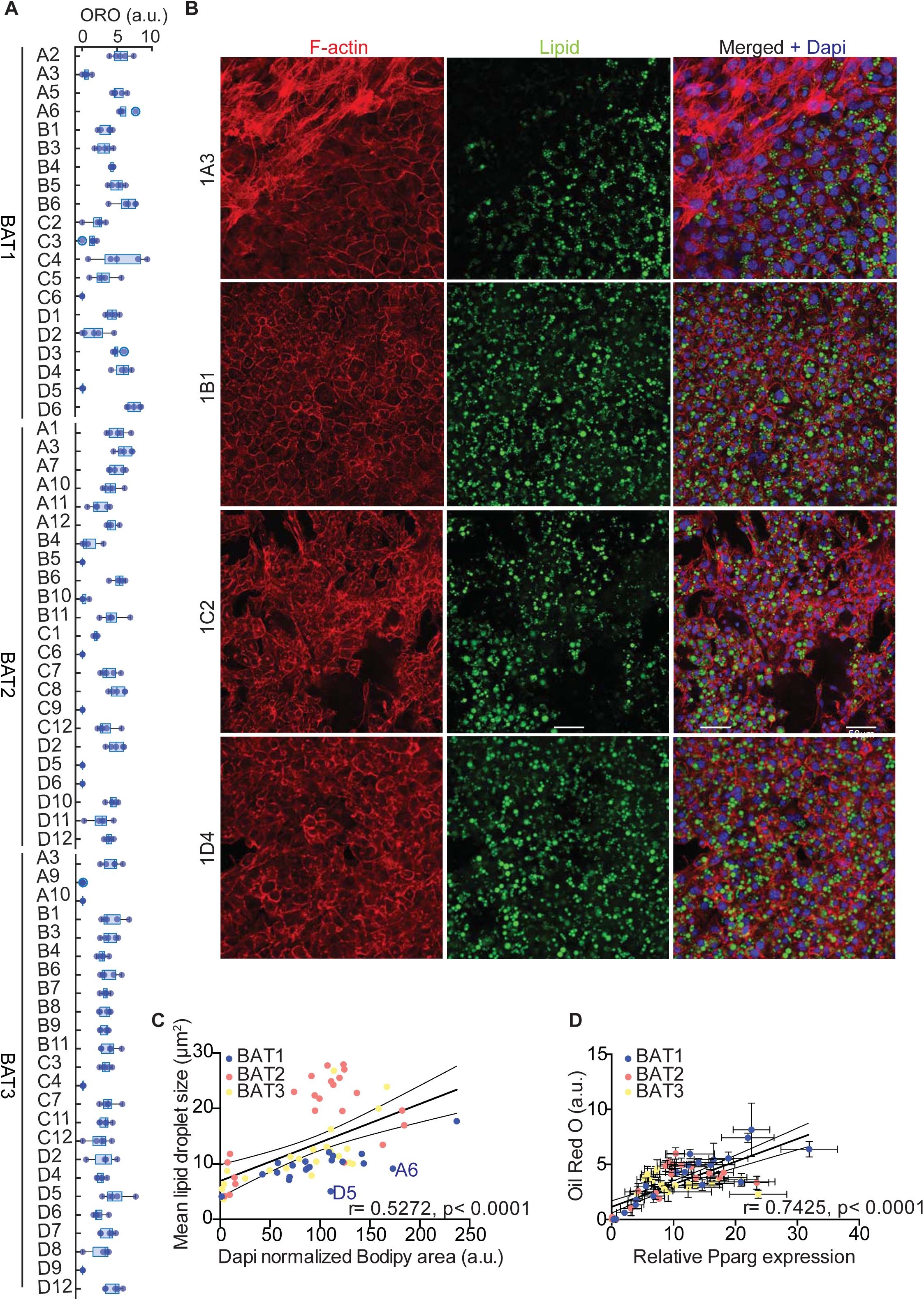
Heterogeneity in differentiation capacity and lipid accumulation of brown adipocyte clones. (**A**) Quantification of relative lipid accumulation measured by ORO staining at day 8 of differentiation from 67 immortalized brown preadipocyte clones (n= 5; mean OD normalized to DAPI ± SEM). (**B**) Representative images of differentiated cell lines, stained with F-actin (red), lipid droplets (green), and DAPI (blue). (**C**) Correlation between mean lipid droplet size and lipid area normalized by the number of nuclei per clone (n> 200), calculated from pictures shown in Supplementary Data 2. Values are mean of different area scanned per clones (n= 9). (**D**) Correlation analysis of Pparg expression and lipid accumulation (n=5).

**Figure 3:**
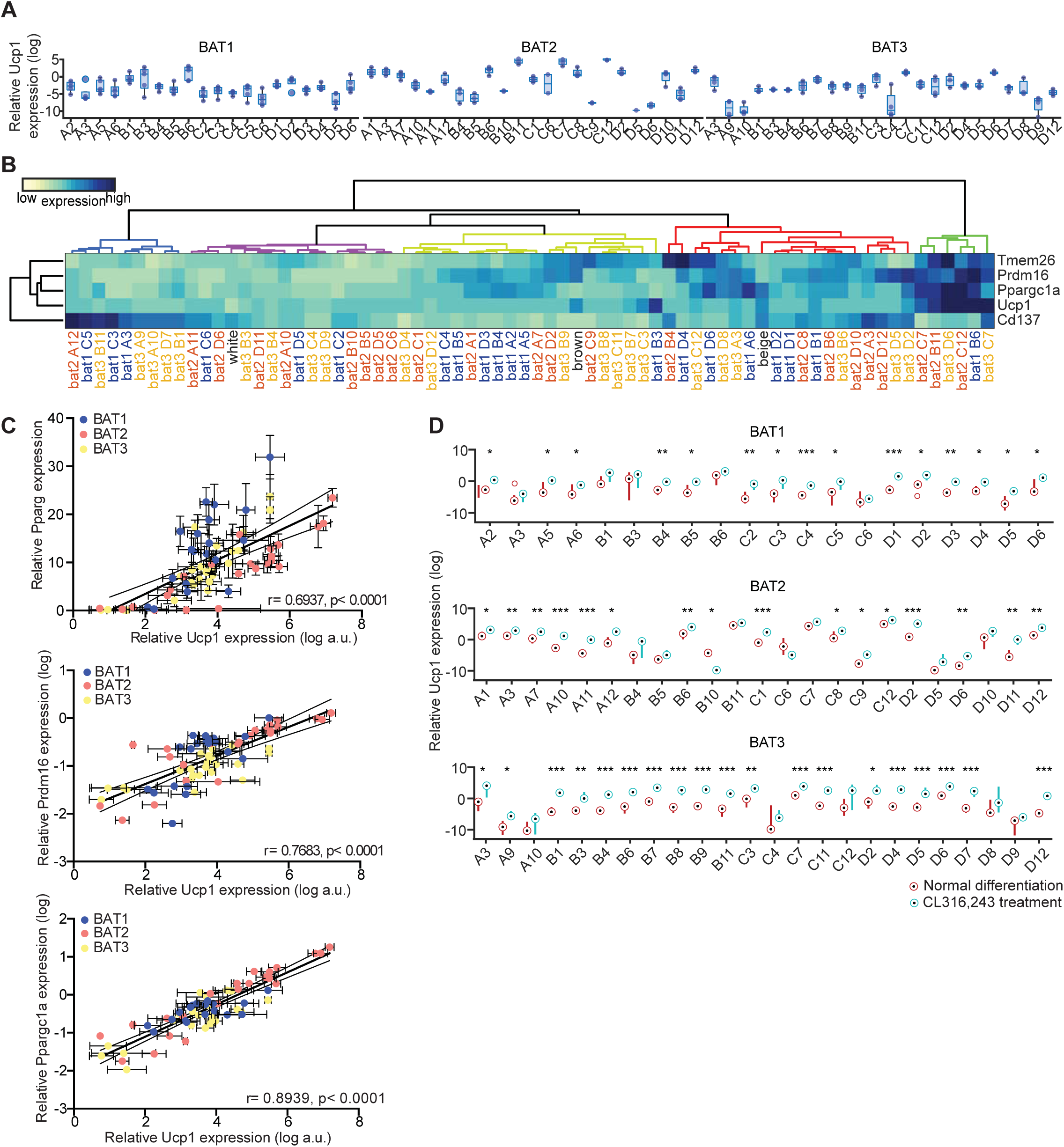
Brown adipocyte clones heterogeneously express brown, beige and white fat markers. (**A**) mRNA expressions of Ucp1 at day 8 of differentiation (n=4-5). (**B**) Heatmap of mRNA expression for different adipocyte markers in all 67 clones, as well as in vitro differentiated white, beige and brown adipocytes at day 8 of differentiation (n=3-5). Median expression for each cell line was transformed to log scale and gene wise z-scores were computed independently for each cell line. Dendogram colors denote identified adipocyte clone cluster. (**C**) Correlation of Ucp1 with Pparg (left panel), Prdm16 (middle panel) and Ppargc1a (right panel) expressions. The values were mean ± SEM, and log transformed for Ucp1, Prdm16 and Ppargc1a (n= 4-5). Gene expression was normalized to TBP. (**D**) Ucp1 expression of controls and cells treated for 3 hours with 0.5μM CL-316,243 (n=3-5).

Next, we tested expression of additional markers for brown (Prdm16, Ppargc1a) and beige (Cd137, Tmem26) adipocytes (*17, 34*) and performed unsupervised hierarchical clustering (Euclidean distance and Ward’s linkage method) of the data to see if the clonal cell lines could be grouped using these markers. For comparison, we also included *in vitro* differentiated brown, beige and subcutaneous white adipocytes in the analysis (**Fig. 3B** and **Supplementary Fig. 2**). 19 cell lines clustered with brown adipocytes, 18 with beige and 15 with white adipocytes, with two additional clusters containing 6 and 9 cell lines with either very high or very low thermogenic gene expression, respectively. The latter cluster was largely defined as Cd137^high^ and low for all other markers (**Fig. 3B**). The 67 different cell lines expressed very different combinations of brown, beige and white adipocyte markers. Expression of Prdm16 and Ppargc1a, two key factors determining differentiation of preadipocytes towards brown adipocytes (*9, 35, 36*) were largely co-expressed with Ucp1 (**Fig. 3B**). This was also observed in a correlation between Prdm16, Ppargc1a and Ucp1, respectively, while the correlation between Pparg and Ucp1 expression was weaker (**Fig. 3C**). However, there were several clones that revealed a surprising dissociation of expression between these markers for brown adipocytes. The beige adipocyte markers Cd137 and Tmem26 were not expressed in the same pattern, with highest expression of these genes in different non-overlapping cell lines. Interestingly, inter-individual differences were smaller than differences between clonal lines of the same mouse. This suggests that there is developmental heterogeneity between preadipocytes. Based on the observed differences in brown, beige and white characteristics of the clonal cell lines, we tested cellular response to acute (0.5 μM for 3 hours) β3-adrenergic receptor agonist (CL-316,243) treatment, with respect to Ucp1 gene expression (**Fig. 3D**). Ucp1 expression was induced in most cell lines by acute CL-316,243 treatment, except 1A3, 1C6, 2B4, 2B5, 2B10, 2C6, 2D5, 3A10, 3C4 and 3D9, which were also the least differentiated cell lines (**Fig. 3D and 2A**). Thus, the data obtained from these 67 clonal brown adipocyte cell lines strongly suggested functional and developmental heterogeneity.

### RNAseq expression profiling reveals differences in brown preadipocytes and adipocytes

To obtain a more detailed view on the transcriptional differences of the cell lines, we performed RNA sequencing (RNAseq) from all 20 undifferentiated and differentiated cell lines of mouse 1, that all showed comparable proliferation rates (**Supplementary Fig. 3A**), indicating no differences in the effects of SV40 immortalization. After pre-processing and filtering (*37*) a total of 9483 for undifferentiated and 10363 genes for differentiated brown adipocytes remained for further analysis.

Unsupervised hierarchical clustering for pre- and differentiated adipocytes did not reveal any common pattern of conserved cell identities between undifferentiated and differentiated cells (**Supplementary Fig. 3B**). This was also confirmed by PCA, which did not indicate any conclusive clustering or pattern that would allow for the identification of distinct brown adipocyte lineages (**Supplementary Fig. 3C**). To assess how our dataset compared to gene expression profiles of whole white and brown adipose tissues and different cell lines, we analyzed our data sets using the ProFAT database (*38*). This online tool is designed to provide a relative estimation of the brown adipose tissue characteristics of the analyzed samples (*38*). Since all cell lines were derived from BAT, we display the data by similarity to brown fat “BATness”, scaled from 0 (low) to 1 (high). We performed pairwise comparisons for each undifferentiated and differentiated sample of each cell line (**Supplementary Fig. 3D**). Using k-means clustering, we identified three different clusters. One cluster did not show characteristics of BAT in either the undifferentiated or differentiated states (dots); the second acquired brown fat characteristics upon differentiation (triangles); whereas the third one showed BAT characteristics in both undifferentiated and differentiated cells (squares) (**Supplementary Fig. 3D**).

### Laplacian Eigenmap based dimensionality reduction reveals distinct brown adipocyte gene expression signatures

This classification indicated that there are groups of cell lines with distinct and shared molecular characteristics. However, the ProFAT associated clusters did not correlate with clusters in the hierarchical clustering of the RNAseq data. Furthermore, based on the previous analysis we were neither able to estimate the number of brown adipocyte lineages among our clones, nor could we identify marker genes (features) allowing us to group the cell lines into lineages.

Nevertheless, the RNA expression data and ProFAT correlations also suggested cellular heterogeneity in murine BAT. Consequently, we applied nonlinear dimensionality reduction techniques to uncover hidden patterns aiming to further classify our cell lines. To avoid a bias based on differences in differentiation capacity, and to define distinct precursor populations, we restricted our analysis to the RNAseq data from preadipocytes. After applying Laplacian Eigenmaps, the projected gene expression revealed seven distinct gene sets, from which we generated seven gene expression modules (GEMs) M1-M7, with sizes between 76 and 227 genes (**Fig. 4A, Supplementary Table 2**).

**Figure 4:**
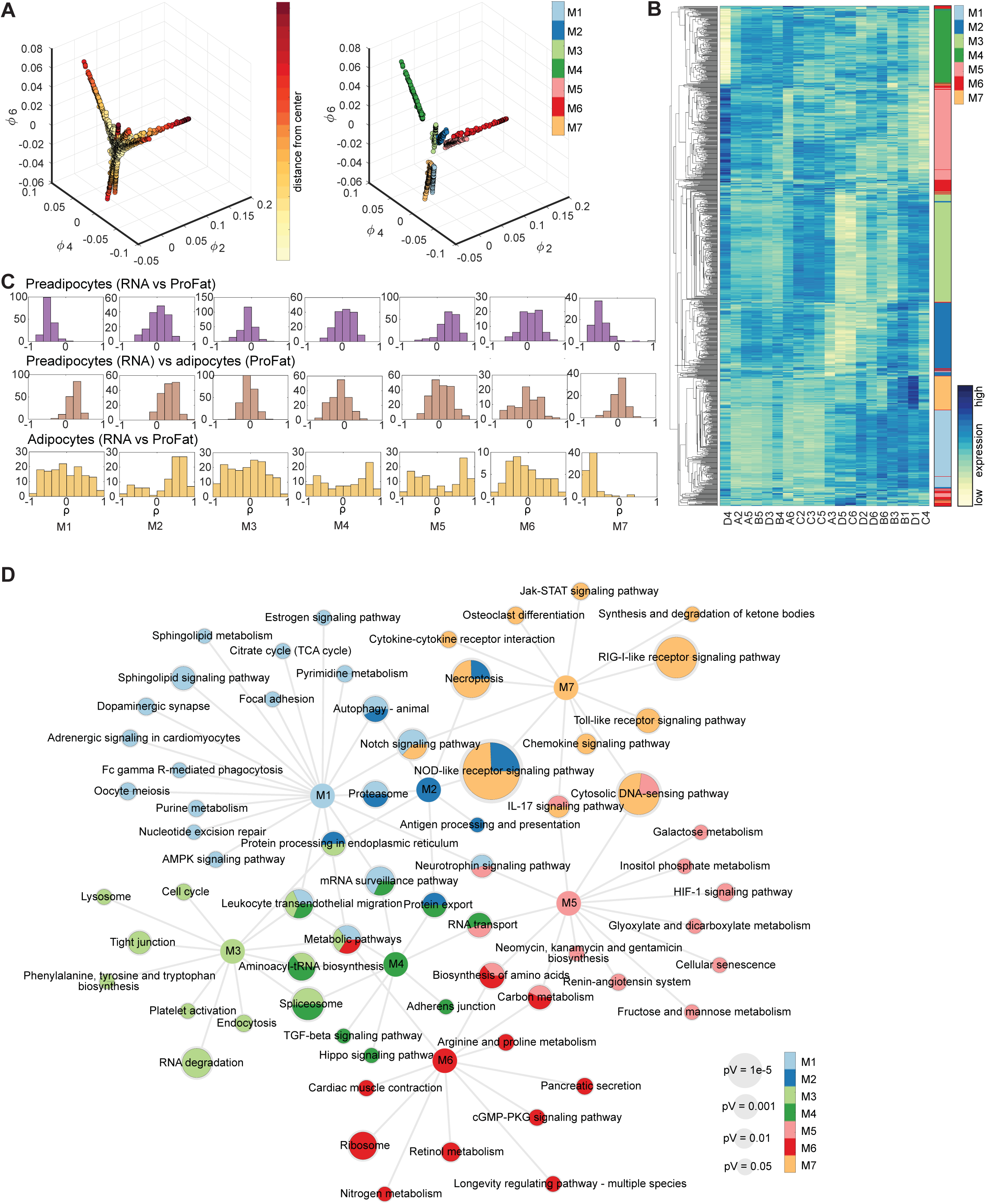
Laplacian Eigenmap based extraction of expression modules. (**A**) Brown preadipocyte gene expression in Laplacian Eigenmap derived low dimensional space. Expression of the three eigenvectors ϕ2, ϕ4 and ϕ6. Distance from estimated center is color-coded (left panel). After removal of genes with low variance module membership is coded in colors (right panel). (**B**) Heatmap of hierarchical clustered module genes in rows. Gene expression was z-score normalized. Module membership is indicated by the right color bar. (**C**) Distribution of correlation coefficients of the gene wise comparison of expression to estimated BATness for each module. Upper line: pre-adipocyte gene expressions vs estimated BATness of pre-adipocytes from ProFat database. Middle line: pre-adipocyte expressions vs differentiated BATness from Profat database. Lower line: differentiated adipocyte expressions vs differentiated BATness from ProFat database. (**D**) Network of KEGG enriched pathways. Significantly enriched pathways are connected to the respective module nodes. Size of the Pathway nodes and proportion of the node-pie-chart refer to –log10 of enrichment p-value. Color of the nodes refers to module membership.

We next analyzed if the genes defining the seven GEMs allow defining distinct cellular lineages or populations. Almost all of the 7 GEMs showed distinct gene expression patterns, only module M6 split up in small subgroups distributed all across the clustering tree (**Fig. 4B**). Modules M1-M5 and M7 showed unique expression patterns, thus, each of them allowing for sub classification of the clonal cell lines. However, at this point the data did not support preference of one module over the other. Further, PCA of GEM gene expression of both undifferentiated and differentiated brown adipocyte clones did not reveal clear populations (**Supplementary Fig. 4**). Thus, we compared module specific expression patterns to estimated BATness. We correlated the individual gene expression of each preadipocyte module to the estimated BATness of either the undifferentiated (**Fig. 4C** upper panel) or differentiated (**Fig. 4C** middle panel) cell lines, as well as gene expression and estimated BATness of adipocytes (**Fig. 4C** lower panel) and summarized the correlation coefficients for each gene within the individual modules. In preadipocytes, M1 and M7 contain genes that negatively correlate with BATness, whereas gene expression of M5 is positively correlated to BAT. M2-4 and M6 did not show characteristics that correlate with brown adipocytes. Conversely, genes in M1 and M7 showed a trend to positively correlate with BATness in differentiated adipocytes, similar to M2, whereas all others showed a more random distribution. KEGG enrichment analysis of each GEM showed associations with diverse pathways and little overlap between the modules (**Fig. 4D, Supplementary Table 3**), except M2, which shared most of its enriched pathways except ‘Antigen processing and presentation’ with other modules. M1 showed highest association with sphingolipid signaling and metabolism as well as autophagy and notch signaling, whereas M7, the other WAT associated module was characterized by pathways associated with immune cell signaling. Conversely, M5, with highest BAT correlation, associated with carbohydrate and amino acid metabolism. Taken together, these data showed that the identified GEMs contain information to functionally distinguish our clonal cell lines based on immune signaling, cell adhesion and migration, RNA and amino acid metabolism and fuel processing.

### Eif5, Tcf25 and Bin1 mark subpopulations of brown preadipocytes and adipocytes *in vivo*

To identify potential brown adipocyte lineage markers, we selected those genes with stable expression between preadipocytes and adipocytes. From the seven GEMs, we identified 57 genes showing stable expression (p < 0.05, Pearson correlation) between pre- and differentiated brown adipocyte clones (**Fig. 5A**). A PCA of these genes from either undifferentiated or differentiated brown adipocyte clones also did not clearly separate the cell lines into different clusters (**Supplementary Fig. 5A**). ORO quantification-based PCA on preadipocytes and adipocytes did also not show any obvious clustering of the cell lines (**Supplementary Fig. 5B**). Single gene association to browning was estimated by correlating the gene expressions to cell line specific BATness. Thus, we identified genes that potentially mark cells with distinct characteristics, indicative of distinct lineages (**Fig. 5B**). The two genes from M4 (Bin1 and Phax) were least correlated with BATness, whereas genes from M2 were mainly correlated with BATness. The genes from the other GEMs were more evenly distributed. We selected the eukaryotic translation initiation factor 5 (Eif5), transcription factor 25 (Tcf25) and bridging integrator 1 (Bin1) to test as potential lineage markers from the stably expressed 57 genes, based on BATness, in combination with the maximum expression and fold change in our clonal cell lines as well as the availability of antibodies (**Supplementary Fig. 5C**). Among those three, Eif5 was most highly associated with a classical BAT phenotype; Tcf25 could not be assigned (no tendency to be more-or less-brown); while Bin1 was the least brown adipocyte associated gene (**Fig. 5B**.) We observed a similar pattern when cells were correlated to lipid content (**Supplementary Fig. 5D**). The expression of Eif5, Tcf25 and Bin1 in the differentiated cell lines was correlated with the expression, as assessed by qPCR, of Ucp1, Pparg, Ppargc1a and Prdm16 (**Fig. 5C**). Eif5 was positively correlated with Ucp1, Pparg, Ppargc1a and Prdm16. Tcf25 showed no correlation with Ucp1 and differentiation markers, whereas Bin1 negatively correlated with Ucp1 and the other markers (**Fig. 5C**).

**Figure 5:**
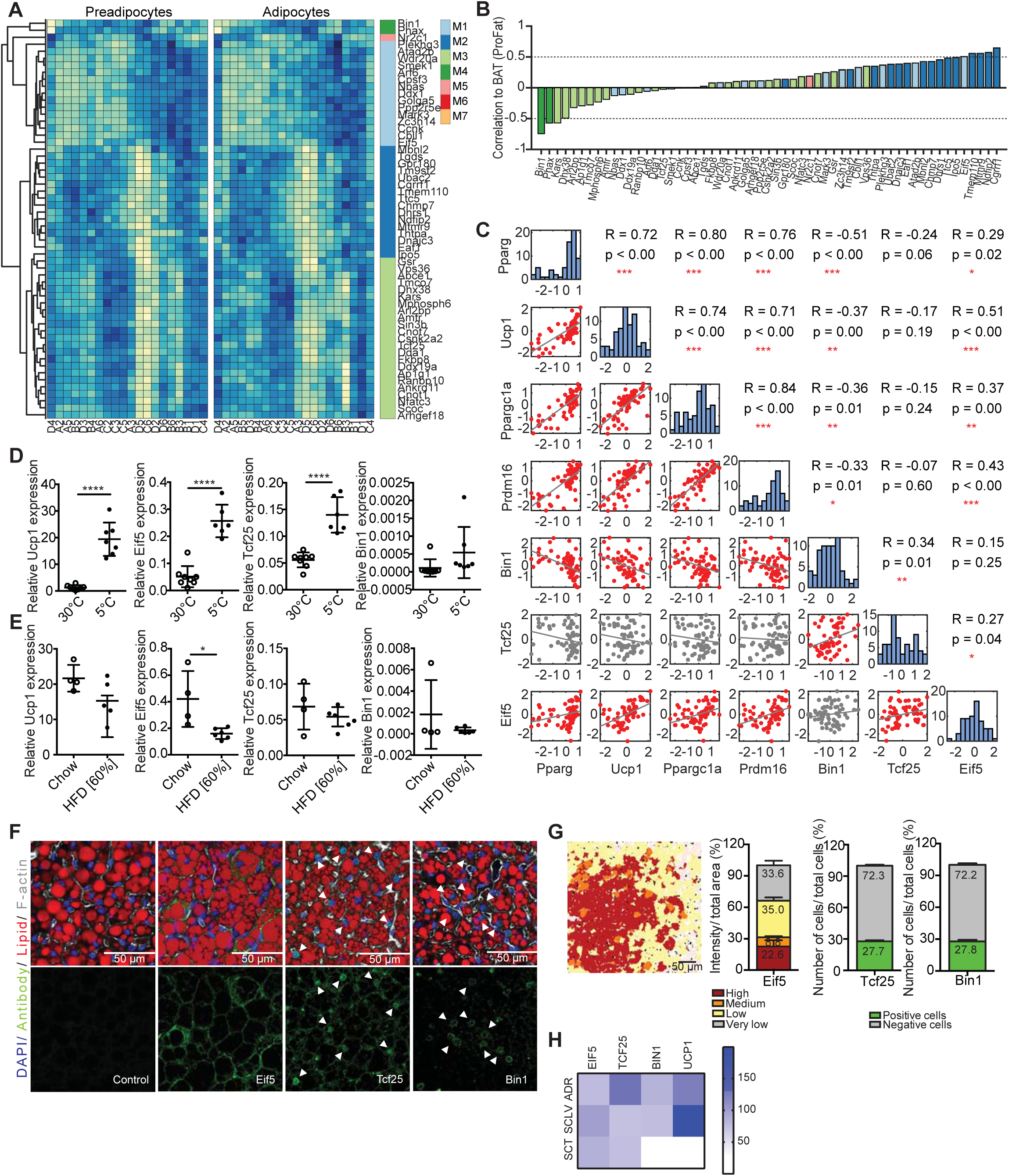
Eif5, Bin1 and Tcf25 mark subsets of brown adipocytes. (**A**) Heatmaps of stably expressed genes of preadipocytes (left panel) and differentiated brown adipocytes (right panel). Module membership is indicated by the right color bar. (**B**) Correlation coefficients for stable expressed genes compared to estimated BATness. Color of bars indicates module membership. (**C**) Correlation plot for pairwise comparison of selected markers to selected stably expressed genes. Red plots denote a significant correlation. (**D**) Expression of Ucp1, Eif5, Tcf25 and Bin1 in mice kept in cold or thermoneutrality (n= 6-8, data are mean expressions normalized to B2m ± SEM). (**E**) Expression analysis of Ucp1, Eif5, Tcf25 and Bin1 in chow- and HFD-fed mice (n= 4-6, data were mean expressions normalized to B2m ± SEM). (**F**) BAT staining for Eif5, Tcf25 or Bin1 (green), F-actin (gray), lipid droplets (red) and DAPI (blue) from wild-type C57BL/6 mice. (**G**) Representative color gradient picture of Eif5 staining on BAT (left panel), and percentage area of different Eif5 intensity (high, medium, low, very low) normalized to total area (second panel, n= 9 sections). Quantification of Tcf25- and Bin1-positive and negative cells in percentage of total cells (n= 9-18 sections). (**H**) Heatmap of mRNA expression in human periadrenal (ADR), supraclavicular (SCLV) and subcutaneous (SCT) adipose tissue from six different donors. A and B based on RNAseq data, remaining analysis based on qPCR data. * p< 0.05, ** p< 0.01, *** p< 0.001, **** p< 0.0001

Next we tested if expression of Eif5, Tcf25 and Bin1 showed similar associations with Ucp1 expression *in vivo*. To this end, we studied gene expression in BAT of mice chronically housed at either thermoneutrality (30°C) or at 5°C (**Fig. 5D**). As previously reported, Ucp1 expression was increased upon chronic cold exposure when compared to its expression at thermoneutrality. Similarly, expression of Eif5 and Tcf25 was increased upon chronic cold exposure, whereas Bin1 expression remained unaltered. Expression of Ucp1 was slightly decreased in BAT of obese mice fed a high fat diet (HFD) compared to lean chow diet animals (**Fig. 5E**). An even more pronounced reduction was observed for Eif5, whereas expression of Tcf25 and Bin1 were not altered by diet-induced obesity (**Fig. 5E**).

Further analysis of our scRNAseq data showed a random distribution of Eif5, Tcf25 and Bin1 in the four preadipocyte clusters, further confirming the stable expression of these markers throughout brown adipocyte differentiation (**Supplementary Fig. 5E**). To investigate the distribution of Eif5, Tcf25 and Bin1 in mature adipocytes *in vivo*, we performed immunostaining for Eif5, Tcf25 and Bin1 on murine BAT sections. Similar to Ucp1, Eif5 expression was detected in most brown adipocytes, with differing intensities (**Fig. 5F** and **G**). Following our quantification strategy applied for Ucp1 (**Fig. 1A**), we quantified Eif5 expression in individual brown adipocytes and grouped the cells according to high, medium, low and very low expression, with around 1/4 of the brown adipocytes showing high or very high expression (**Fig. 5G, Supplementary Fig. 5F**). Nuclear or perinuclear expression of Tcf25 or Bin1, respectively was observed in ∼ 25% of brown adipocytes (**Fig. 5F** and **G**). In addition, Bin1 staining was also observed in endothelial cells. Comparative gene expression analysis in periadrenal, supraclavicular and subcutaneous white and brown adipose tissue of humans confirmed expression of EIF5, TCF25 and BIN1 in human BAT, but also in WAT as seen in mice (**Fig. 5H, Supplementary Fig. 5G** and **H**).

### Loss of Bin1 increases Ucp1 expression and oxygen consumption

To functionally characterize differences of Eif5^high^, Tcf25^high^ and Bin1^high^ brown adipocytes, we selected representative clones [B1 (Eif5^high^), D1 (Tcf25^high^) and D5 (Bin1^high^)] using hierarchical clustering from gene expressions measured by qPCR (**Fig. 6A**). We established stable knockdown cell lines of these genes in the respective clones, which was confirmed by semi-quantitative PCR of respective gene expression on day 0 (preadipocytes) and day 8 (adipocytes) of differentiation (**Fig. 6B**). Neither of the knockdowns impaired adipogenesis as assessed by expression of Pparγ (**Supplementary Fig. 6A**). However, reduction of Eif5 in the B1 clone induced Ucp1 and P2rx5 mRNA expression (**Supplementary Fig. 6A**), but had no significant effect on mitochondrial respiratory capacity (**Fig. 6C**, left panel). Tcf25 knockdown in clone D1 reduced expression of Ucp1 and Prdm16 and showed increased expression of Cd137 and the white adipocyte marker Asc1 (*16*) in comparison to its shScr control (**Supplementary Fig. 6A**). Microplate-based oxygen consumption of the shTcf25 D1 clone revealed reduced maximal respiratory capacity (**Fig. 6C**, middle panel), but no differences in mitochondrial uncoupling. Knockdown of Bin1 in the D5 clone resulted in an up-regulation of Ucp1 mRNA levels (**Supplementary Fig. 6A**) and showed a higher basal oxygen consumption rate (OCR). Addition of oligomycin (ATP synthase inhibitor) to the wells did not reduce the OCR of shBin1 D5, indicating full mitochondrial uncoupling (**Fig. 6C**, right panel). Thus, expression of Bin1 in preadipocytes and adipocytes correlates with low thermogenic capacity, and loss of Bin1 results in increased thermogenic gene expression and full mitochondrial uncoupling, at least in clones with high Bin1 expression.

**Figure 6:**
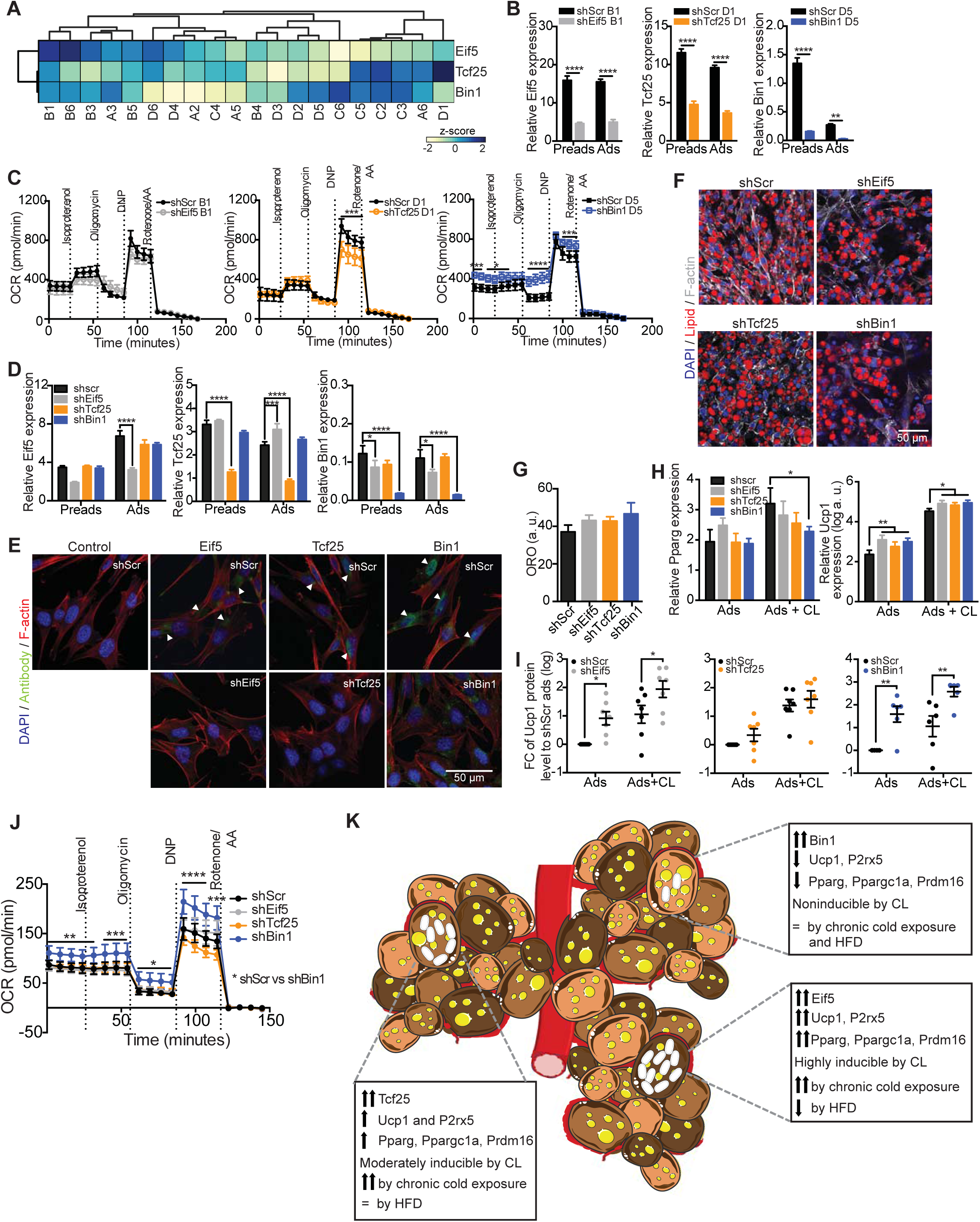
Loss of Bin1 increases basal respiration and UCP1 protein content. (**A**) Heatmap and hierarchical clustering of cell lines with regards to the mRNA expressions of Eif5, Tcf25 and Bin1. (**B**) Expression of Eif5, Tcf25 and Bin1 in control cells (shScr) of different clones (B1, D1 and D5) and the respective knockdown (shEif5 B1, shTcf25 D1 and shBin1 D5); Preads (d0 of differentiation) and Ads (d8 of differentiation) (n=5). (**C**) OCR of shScr B1 and shEif5 B1 (left panel), shScr D1 and shTcf25 D1 (middle panel) and shScr D5 and shBin1 D5 (right panel) at day 8 of differentiation measured by Seahorse (n=4). (**D**) Expression of Eif5, Tcf25 and Bin1 in control cells (shScr) and respective knock-down (shEif5, shTcf25 and shBin1); Preads (d0 of differentiation) and Ads (d8 of differentiation) (n=5). (**E**) Immunofluorescence staining of Eif5/Tcf25/Bin1 (green), F-actin (red) and DAPI (blue) on shScr/shEif5/shTcf25/shBin1 preadipocytes. (**F**) Representative images of differentiated cell lines (shScr, shEif5, shTcf25 and shBin1) stained with F-actin (white), lipid droplets (red), and DAPI (blue). (**G**) Quantification of relative lipid accumulation measured by ORO staining at day 8 of differentiation from knockdown cell lines (n=3, mean OD normalized to DAPI ± SEM). (**H**) mRNA expressions of Pparg (left panel) and Ucp1 (right panel) on shScr, shEif5, shTcf25 and shBin1 adipocytes and CL treated adipocytes (0.5μM CL-316,243 for 3 hours; n=6). (**I**) Quantification of western blots for Ucp1 (Abcam, cat# 10983) in shScr, shEif5, shTcf25 and shBin1 adipocytes and adipocytes treated with 0.5μM CL-316,243 for 6 hours (n=6-7) normalized to β-actin. Values are fold changes to the respective shScr adipocytes (ads). (**J**) OCR of control and knockdown cell lines at day 8 of differentiation measured by Seahorse (n=3); data were normalized by non-mitochondrial respiration. (**K**) Schematic illustration of brown adipose tissue heterogeneity: Murine brown adipose tissue is composed of functionally distinct brown adipocytes. We identify Eif5 expressing brown adipocytes as “classical” brown adipocytes, with high UCP-1 content and mitochondrial uncoupling. Tcf25 expressing brown adipocytes are similar, albeit with lower UCP-1 expression. In contrast, Bin1 expressing brown adipocytes appear in a dormant state, expressing low UCP-1 levels with little response to beta adrenergic stimulation. However, loss of Bin1 or chemical mitochondrial uncoupling reveals high thermogenic capacity of these cells. Subpopulations of brown adipocytes are color coded regarding to their thermogenic capacity, with the highest shown in dark brown (Eif5^high^), followed by brown (Tcf25^high^) and light brown (Bin1^high^) (= indicates unchanged expression). All qPCR data were normalized to TBP and shown as mean ± SEM. * p< 0.05, **, ^##^ p< 0.01, *** p< 0.001, ****p< 0.0001

We next tested the effects of stable depletion of these markers in a mixed brown preadipocyte population. To this end, we created brown preadipocyte knockdown cell lines for Eif5 (shEif5), Tcf25 (shTcf25) or Bin1 (shBin1), with scrambled shRNA expressing cells (shScr) as the control. qPCR analysis in preadipocytes and fully differentiated adipocytes confirmed a stable knocked-down of all three genes (**Fig. 6D**). This was further confirmed by immunofluorescence stainings on preadipocytes (**Fig. 6D**). As with the clonal cell lines we did not observe any differences in differentiation capacity upon loss of either of the marker genes with respect to lipid accumulation (**Fig. 6F and G)** or expression of Pparg (**Fig. 6H**, left panel), despite a reduction in Pparg expression in CL-316,243 treated shBin1 cells (**Fig. 6H**). Cd137 mRNA levels were generally higher in the knockdown adipocytes compared to controls, whereas Prdm16 expression was lower in shEif5 and shTcf25 lines compared to control cells (**Supplementary Fig. 6B**). In line with the clonal knockdown data, depletion of Tcf25 resulted in increased expression of the white adipocyte marker Asc-1 and a downregulation of the brown adipocyte marker P2RX5 (**Supplementary Fig. 6B**). In contrast, shEif5 and shBin1 cells had significantly higher mRNA or protein levels of Ucp1 without or with CL 316,243 treatment compared to the shScr adipocytes, respectively (**Fig. 6H** and **I**). shBin1 cells also expressed highest levels of P2rx5 (**Supplementary Fig. 6B**). Seahorse analysis revealed that oxygen consumption was highest in shBin1 cells (**Fig. 6J**). Thus, Bin1 marks a relatively “white adipocyte like” brown adipocyte population. Moreover, loss of Bin1 in brown adipocytes increases thermogenic gene expression and mitochondrial activity suggesting that Bin1 functions in suppressing classical brown adipocyte identity.

Taken together, these data demonstrate that using our combined approach we were able to identify a set of genes, including Eif5, Tcf25 and Bin1, with preserved expression between preadipocytes and adipocytes that mark subsets of functionally distinct brown adipocytes within BAT. Our data support a model where Bin1 suppresses full mitochondrial uncoupling; marking a “dormant” brown adipocyte subpopulation, whereas Eif5 and Tcf25 positive brown adipocytes represent “classical” fully uncoupled brown adipocytes. Thus, our data strongly suggest the presence of multiple brown adipocyte lineages in mice (**Fig. 6K**).

## Discussion

In the current study we aimed to clarify the presence of distinct brown adipocyte lineages in mice. We find that even at room temperature, where murine BAT appears morphologically uniform, Ucp1 expression within individual brown adipocytes differs greatly throughout the tissue. To this end, we tested if differences in Ucp1 content are the result of developmental heterogeneity among murine brown adipocytes and not due to anatomical or environmental differences. Using scRNAseq from the SVF of adult murine BAT, we identified a diverse set of immune cell and preadipocyte clusters. Surprisingly, a more detailed analysis of the preadipocyte clusters indicated that clustering within preadipocytes was driven by differentiation rather than distinct developmental lineages. Therefore, albeit providing an interesting resource for others to study cellular composition within BAT, these scRNAseq data did not allow identification of distinct brown preadipocyte lineages. To overcome this limitation we generated, similar to a previous study on human brown adipocyte cell lines (*39*), 67 immortalized brown preadipocyte clones from lean adult male C57Bl/6 mice housed at room temperature. Thus, as currently believed, all preadipocytes isolated from the BAT of this mouse should, in contrast to precursors isolated from human BAT (*39*), by definition give rise to only brown adipocytes. We show that these clones differentiate, albeit to various degrees, into lipid laden adipocytes, expressing adipogenic markers such as Pparg. However, these cell lines were heterogeneous with respect to Ucp1 expression, which was not well correlated with the capacity to differentiate and accumulate lipids.

Heterogeneity between the cell lines was also apparent when comparing their response to β3-adrenergic stimulation or the expression of classical brown, beige and white adipocyte markers. Thus, similar to Shinoda et al, we observe strong differences in gene expression between the clonal cell lines (*39*).

To obtain more insights into the differences between the clonal cell lines, we focused on the clones from one mouse and performed RNAseq from all undifferentiated and differentiated cell lines. Selecting clones from one donor is critical to exclude studying inter-individual differences rather than differences between individual cellular lineages. Neither clustering nor PCA analysis of the data showed a conserved structure between preadipocytes and adipocytes that would allow us to group the cell lines into lineages. This suggests as also seen in the scRNAseq data that there are many subtle differences between the clonal cell lines, influenced by cell cycle state, proliferation capacity etc., which generate transcriptional noise complicating the assignment of individual cell lines into lineages.

Hence, we compared the RNAseq profile of each preadipocyte and adipocyte sample to the ProFat database to assess for each cell line its relative browning capacity. By doing so we were able to identify three clusters of cells that showed characteristics of BAT in both the undifferentiated and differentiated states, acquired brown fat characteristics upon differentiation, or showed no BAT characteristics in both undifferentiated and differentiated cells. This comparison clearly demonstrated a separation of our cell lines, albeit all derived from the SVF of BAT, strongly indicating distinct lineages of our clones.

To identify lineage markers that we could use to visualize these clusters/lineages *in vivo*, we used Laplacian Eigenmaps to structure our data identifying seven distinct gene expression modules. From these genes we selected those with conserved expression between preadipocytes and mature adipocytes and correlated each of them with the ProFat database. From these 57 genes we selected Eif5, Tcf25 and Bin1 as representatives for brown adipocytes with a classical brown adipocyte, intermediate or more distant gene expression, respectively. We substantiated our *in vitro* findings through stainings of mature brown adipocytes in BAT for Eif5, Tcf25 and Bin1 revealing a contribution of each population to around 25% of brown adipocytes.

Moreover, it is important to note that in contrast to our previous work identifying white, beige and brown adipocyte specific surface proteins (*16*), neither of these genes is brown adipose selective. In contrast, all three genes show a broad tissue expression pattern. However, using different environmental challenges such as chronic cold exposure and HFD feeding, we find that Eif5 and Tcf25 expression closely parallel the expression of Ucp1, whereas Bin1 expression was not regulated under each of these conditions in BAT *in vivo* (**Fig. 6K**).

The eukaryotic translation initiation factor 5 (Eif5), functions to initiate protein synthesis through interaction with the 40S ribosomal subunit (*40, 41*). Transcription factor 25 (Tcf25) has been shown to play an important role during early embryonic organogenesis (*42*) and its expression is decreased by age in several organs, such as kidney, heart, liver, and lung (*43*). Bridging integrator 1 (Bin1), or amphiphysin 2, shows the most restricted expression pattern among the three, with highest expression in skeletal muscle and brain and has been shown to regulate muscle differentiation (*44*).

The function of any of these genes in BAT, however, has not been studied yet. We find that loss of Eif5 did not significantly impact on the brown adipocyte characteristics of these cells. Interestingly, loss of Tcf25 shifted cellular identity to a more white-like adipocytes with up-regulation of Asc1, a white adipocyte marker, and reduction of maximum respiratory capacity to respire. On the other hand, Bin1^high^ cells expressed low Ucp1 and Bin1 expression negatively correlated with thermogenic genes. Interestingly, loss of Bin1 resulted in strongly increased Ucp1 expression and mitochondrial activity, indicating that Bin1 is not only a marker of “quiescent” brown adipocytes, but actively suppresses thermogenic capacity (**Fig. 6k**).

We also show that the three markers Eif5, Tcf25 and Bin1 are present in human BAT, strongly supporting the notion that distinct brown adipocyte populations could also exist in humans. Yet it is important to note that there may be several other lineage markers to be identified that will help create an overall picture of interspecies brown adipocyte lineage determination and functional development. Moreover, future studies on adipose selective deletion of these putative lineage markers will need to address the specific function of these proteins in brown fat.

Nevertheless, activation of brown adipose tissue remains an attractive pharmacological goal to increase energy expenditure and combat the ever growing pandemic of obesity and the metabolic syndrome (*45-47*). However, anatomical, morphological and potentially functional and developmental differences between rodent and human BAT complicate the translation from rodents to human physiology (*26*). At room temperature, morphology of murine BAT appears homogeneous, while human BAT appears as a mixture of unilocular and multilocular adipocytes (*48, 49*).

Conversely, at thermoneutrality or upon prolonged high fat diet feeding, murine BAT resembles the morphology of human BAT (*49, 50*). The appearance of unilocular adipocytes could either indicate the *de novo* differentiation of white adipocytes or excessive accumulation of triglycerides in brown adipocytes. The latter would suggest that individual brown adipocytes within murine BAT respond very differently to environmental changes, such as temperature and diet and this could indicate the existence of distinct brown adipocyte lineages with potentially different function. Moreover, if BAT were also composed of multiple brown adipocyte lineages in humans, differences in the relative contribution of these lineages could help explain differences between individuals with respect to BAT activity and the difficulties to translate findings from rodents housed at room temperature to humans.

In conclusion, we provide the first evidence for the existence of at least three functionally distinct brown adipocyte lineages in mice that differ in their activation state (mitochondrial uncoupling and UCP-1 expression). These data will foster translation by providing novel approaches to “humanize” murine BAT. Moreover, the presence of multiple human brown adipocyte lineages and potential differences in the cellular composition between individuals or upon obesity and the metabolic syndrome could provide novel strategies for personalized obesity therapy.

## Methods

### Cell models and cell culture

Brown preadipocytes were isolated from interscapular BAT of three 8 week old, male WT C57BL/6 mice. BAT was minced and digested with 1mg/ml type-A collagenase (Gibco) in DMEM (Gibco) for 30 minutes at 37°C. The stromal vascular fraction (SVF), was isolated by centrifugation, plated on cell culture plates and subsequently immortalized using an ecotropic retrovirus encoding the SV-40 large T antigen. Single cell clones were picked, following a low density plating of the immortalized preadipocytes and grown individually in separated plates.

Brown preadipocyte clones were cultured in normal growth medium (DMEM + GlutaMAX™, 4.5 g/L D-glucose, pyruvate, 10% FBS and 1% Pen-Strep). Differentiation of preadipocytes was induced with 0.5 mM IBMX, 5 μM dexamethasone, 125 μM indomethacin, 1 nM T3, and 100 nM human insulin, in growth medium for 2 days. Subsequently, medium was changed every 2 days with growth medium containing 1 nM T3 and 100 nM insulin, and cells were harvested at day 8. For acute CL 316,243 treatment, cells were stimulated with 0.5 μM CL for 3 hours, and chronically with 0.1 μM CL 316,243 between days 2-8. Cell cultures were tested regularly negative for mycoplasma contamination.

Brown and subcutaneous preadipocytes were isolated and immortalized from an 8 week old male WT C57/BL6 mouse following the same protocol as above. Mature brown adipocytes were harvested at day 8 of differentiation as stated before. Subcutaneous preadipocytes were differentiated into white and beige adipocytes. For white adipocytes, subcutaneous preadipocytes were induced with 0.5 mM IBMX, 5 μM dexamethasone and 100 nM human insulin in growth medium for 2 days. In addition, 1 μM Rosiglitazone was supplemented to the same induction mix in growth medium for 2 days to differentiate subcutaneous preadipocytes into beige adipocytes. Medium was changed every 2 days with growth medium containing 100 nM insulin. Beige and white adipocytes were harvested at day 8 of differentiation.

For knockdown experiments, immortalized brown preadipocytes were infected with ecotropic lentiviral particles (Cell Biolabs, Inc.) with a scrambled (Scr), anti-Eif5 shRNA (TRCN0000313045: CCGGGACGTTGCAAAGGCGCTTAATCTCGAGATTAAGCGCCTTTGCAACGTC TTTTTG), anti-Tcf25 shRNA (TRCN0000241031: CCGGTGGAAAGAACCCGCCATTATGCTCGAGCATAATGGCGGGTTCTTTCCA TTTTTG), or anti-Bin1 shRNA (TRCN0000238138: CCGGCCCGAGTGTGAAGAACCTTTCCTCGAGGAAAGGTTCTTCACACTCGGG TTTTTG). After 24 hours, cells were selected with growth medium containing 2.5 μg/mL puromycin (Biomol). Cells were maintained and cultured in growth medium with puromycin. CL 316,243 treatment on all knockdown experiments was performed by adding 0.5 μM CL into growth medium containing 1 nM T3 and 100 nM insulin on day 8 of differentiation. Cells were harvested 6 hours after the treatment, followed by RNA or protein extraction.

### Animal models

All animal studies were performed in a conventional animal facility with *ad libitum* access to food and water, 45-65% humidity, and a 12h light-dark cycle. For the HFD study, wild-type male C57BL/6NCrl mice were purchased from Charles River (Germany) at the age of 6 weeks and maintained at constant ambient temperature of 22± 2°C. The mice were fed with low-fat containing control diet (Ssniff E15745-00). At the age of 7 weeks, mice were either maintained on the control diet or were switched to a 60% kcal from fat HFD (Ssniff E15742-34) for 12 weeks. The mice were sacrificed with Ketamine (100mg/kg)/Xylazine (7mg/kg); the interscapular BAT was excised and frozen on dry ice.

Immunofluorescence pictures of Ucp1-KO BAT were taken from Ucp1-KO mice on a C57BL/6J (Jackson Laboratory) genetic background (strain name: B6.129-UCP1tm1Kz/J). The mice were bred, born and weaned at 30°C. With 3 weeks of age the animals were transferred to room temperature and euthanized after 45 days. For the cold exposure study, C57BL/6J mice were bred, born and raised at 30°C. At the age of 10-12 weeks, mice were single housed and randomly assigned to thermoneutral conditions (30°C) or cold (2-3 weeks at 18°C followed by 4 weeks at 5°C) acclimation. Afterwards, mice were euthanized 3-4 hours after lights went on, and BAT was collected. Animal experiments were conducted under permission and according to the German animal welfare law, relevant guidelines and regulations of the government of Upper Bavaria (Bavaria, Germany).

### Lipid staining

Mature brown adipocytes were fixed in 10% formaldehyde for 1 hour at room temperature and dehydrated with 60% isopropanol. Cells were stained with 60% oil Red O in water [stock: 0.35% Oil Red O (Sigma cat# O-0625) in 100% isopropanol] for 10 minutes and subsequently washed six times with ddH_2_O. DAPI staining was used for normalization, and the fluorescence signal was measured at 460 nm, using a microplate reader (PHERAstar FSX). To quantify ORO content, ORO was eluted with 100% isopropanol and measured at 505 nm (PHERAstar FSX).

### MTT assay

Preadipocyte clones were seeded in 96 well plates (2000 cells/well). MTT stock solution was prepared by diluting 50 mg MTT powder (Serva) in10 mL sterile PBS. On the day of measurement, 200 µL MTT stock solution were diluted in 1.6 ml DMEM and added to the wells. As the death control, 200 µL of DMEM medium containing 0.003% Triton X-100 and 0.05% MTT was added to the wells. After 2 hours medium was removed and replaced by 100 µL solubilization buffer (10% Triton X-100 and 0.03% HCl in 100% isopropanol). The plate was incubated for 10 minutes at 24°C at 700 rpm. The lysate was then transferred to a new transparent 96 well plate and measured at 570 nm (PHERAstar FSX).

### qPCR and (sc)RNAseq

RNA from mature brown adipocytes was extracted using the QuickExtract™ RNA extraction kit (Biozym), following the manufacturer’s instructions. cDNA was synthesized by High-Capacity cDNA Reverse Transcription Kit (Applided Biosystem™). Semiquantitative qPCR was performed using iTaq™ Universal SYBR^®^ Green Supermix (Bio-Rad) in a CFX384 Touch Real-Time PCR Detection System (Bio-Rad). Relative mRNA expression was calculated after normalization to TATA-binding protein (Tbp) for cells, and β2-microglobulin (B2m) for tissues unless indicated otherwise in the figure legend.

For human gene expression analysis, tissue from 6 autopsies conducted at the NIH Clinical Center was collected from the following anatomic sites: superficial subcutaneous fat from the anterior abdomen, deep supraclavicular fat and periadrenal fat. Resected tissue was rinsed in PBS and placed immediately in RNAlater (Qiagen). RNA was extracted by homogenizing 100µg tissue using an RNeasy Mini Kit (Qiagen) according to the manufacturer’s instructions. Total RNA concentration and purity were determined by spectrophotometer at 260 nm (NanoDrop 2000 UV-Vis Spectrophotometer; Thermo Scientific). RNA (1µg) was converted to cDNA using the High-Capacity cDNA Reverse Transcription Kit (Applied Biosystems). Relative quantification of mRNA was performed with 3.5µL cDNA used in an 11.5µL PCR reaction for ACTB, BIN1, TCF25, EIF5 and UCP1 using SYBR (Bio Basic) for Bin1, Tcf25 and Eif5 and TaqMan Gene Expression Assay for Ucp1. Quantitative RT-PCR assays were run in duplicates and quantified in the ABI Prism 7900 sequence-detection system. All genes were normalized to the expression of the housekeeping gene β-actin. Primers used are listed in **Supplementary Table 1**. For RNAseq, RNA was extracted with an RNeasy kit (Qiagen), following the manufactures instructions. RIN (RNA Integrity Number values were determined using an automated electrophoresis (Agilent 2100 Bioanalyzer) system and only RNAs with a RIN value >9 were used for further processing (**Supplementary Data 2**). Non-strand specific, polyA-enriched RNA sequencing was performed as described earlier (*51*). Briefly, for library preparation, 1 μg of RNA was poly(A) selected, fragmented, and reverse transcribed with the Elute, Prime, Fragment Mix (Illumina). End repair, A-tailing, adaptor ligation, and library enrichment were performed as described in the Low Throughput protocol of the TruSeq RNA Sample Prep Guide (Illumina). RNA libraries were assessed for quality and quantity with the Agilent 2100 BioAnalyzer and the Quant-iT PicoGreen dsDNA Assay Kit (Life Technologies). RNA libraries were sequenced as 100 bp paired-end runs on an Illumina HiSeq4000 platform. The STAR aligner (*52*) (v 2.4.2a) with modified parameter settings (--twopassMode=Basic) is used for split-read alignment against the human genome assembly mm9 (NCBI37) and UCSC knownGene annotation. To quantify the number of reads mapping to annotated genes we use HTseq-count (*53*) (v0.6.0). FPKM (Fragments Per Kilobase of transcript per Million fragments mapped) values are calculated using custom scripts. A sample overview is provided in **Supplementary Table 4**.

#### scRNAseq

Single cell libraries were generated using the ChromiumTM Single cell 3’library and gel bead kit v2 (PN #120237) from 10x Genomics. Briefly, live cells from the stromal vascular fraction of brown adipose tissue of eight weeks old mice were obtained by flow cytometry following dead cell exclusion using 7-AAD. Afterwards cells were loaded onto a channel of the 10X chip to produce Gel Bead-in-Emulsions (GEMs). This underwent reverse transcription to barcode RNA before cleanup and cDNA amplification followed by enzymatic fragmentation and 5′adaptor and sample index attachment. Libraries were sequenced on the HiSeq4000 (Illumina) with 150bp paired-end sequencing of read2 and 50,000 reads per cell.

### Western blot

Mature brown adipocytes were lysed with RIPA buffer (50 mM Tris pH 7.4, 150 mM NaCl, 1 mM EDTA, 1% Triton X-100), containing 0.1% SDS, 0.01% protease-inhibitor, 0.01% phosphatase-inhibitor cocktail II, and 0.01% phosphatase-inhibitor cocktail III (all from Sigma). Protein concentrations were measured using a BCA Protein Assay Kit (ThermoFischer Scientific), with BSA dilution series as the standard. Proteins were separated by SDS-PAGE, with Fisher BioReagents™ EZ-Run™ Prestained *Rec* Protein Ladder (ThermoFischer Scientific), as the molecular weight marker and transferred to a PVDF Immobilon-P^SQ^ membrane, 0.2 microns (Merck Millipore). Unspecific binding sites were blocked with 5% BSA/TBS-T. Membranes were incubated with primary antibody solutions, Abcam (cat# ab10983), 1:1000. Membranes were washed three times (each 10 minutes), with 1X PBS before incubation with secondary HRP conjugated antibody (Cell Signaling Technology cat# 7074, 1:10000). β-actin (HRP-linked, Santa Cruz Biotechnology cat# sc-47778, 1:5000) was used as loading control. Quantifications were performed using Image J software.

### Immunostainings

Brown adipocyte clones were cultured on 96 well glass-bottom plates, coated with 0.1% gelatin (Sigma-Aldrich, cat# G1890). Preadipocytes as well as fully differentiated brown adipocytes (day 8) were fixed with 10% formaldehyde and blocked with 3% BSA/PBS. Alexa Fluor^®^ 546 Phalloidin (Invitrogen™ cat# A22283, 150 nM), HCS LipidTOX™ Green Neutral Lipid Stain (Invitrogen™ cat# H34775,1:200), and DAPI (Sigma-Aldrich, 1 μg/mL), were used to stain F-actin, lipids and nuclei, respectively. Cells were imaged using the Operetta High-Content Imaging System (PerkinElmer, 20X magnification). Interscapular BAT was collected from 3-4 months old, male C57BL/6 WT mice, and fixed in 4% paraformaldehyde in PBS for 30 minutes at room temperature on a shaker. BAT was embedded in 4% low melting temperature agarose (Sigma-Aldrich, A9414) in PBS and cut into 70 µm sections using a Leica VT1000 S Vibrating blade microtome. Sections were permeabilized with 1% Triton X-100/PBS on ice for 1 minute, and blocked with 3% BSA/PBS for 1 hour. Primary antibodies were added for detection of Ucp1 (Abcam cat# ab10983, 1:250), Eif5 (Abcam cat# ab170915, 1:100), Tcf25 (Invitrogen cat# PA521418, 1:200) and Bin1 (Abcam cat# ab182562, 1:200), followed by Alexa Fluor^®^ 594 anti-rabbit (Abcam cat# ab150080, 1:400) as secondary antibody, Alexa Fluor™ 647 Phalloidin (Invitrogen™ cat# A22287, 1:100), HCS LipidTOX™ Green Neutral Lipid Stain (Invitrogen™ cat# H34775, 1:200) and DAPI (Sigma-Aldrich, 1:1000). Stained sections were mounted on microscope slides using DAKO fluorescence mounting medium (Agilent cat# S3023). Images were captured using a Leica TCS SP5 Confocal (Leica Microsystems).

Automated quantifications of Ucp1 and Eif5 were performed using the commercially available image analysis software Definiens Developer XD 2 (Definiens AG, Munich, Germany) following a previously published procedure (*54*). A specific ruleset was developed in order to detect and quantify Ucp1 and Eif5 stained tissue: In a first step the images were segmented based on color and shape features. The calculated parameter was the mean staining intensity of Ucp1 or Eif5 in the detected cells and they were divided in four classes using the fluorescent intensity of each marker.

### Seahorse analysis

Cells were plated in 24 or 96 (for knockdown cells) well Seahorse plates and differentiated as described above. The oxygen consumption rate (OCR) was measured at day 8 of differentiation using a XF24 or XF96 Extracellular Flux analyzer (Seahorse). One hour before measurement, cells were equilibrated at 37°C, in assay medium (DMEM D5030 supplemented with 0.2% fatty acid-free BSA, 25 mM glucose, 1 mM sodium pyruvate and 4 mM L-Glutamine (Sigma-Aldrich)). Compounds were diluted in assay medium and loaded to the equilibrated cartridge ports (A: 20 µM isoproterenol, B: 15 µM oligomycin, C: 250 µM dinitrophenol (DNP), D: 40 µM Antimycin A and 25 µM Rotenone) and calibrated. Measurements were taken before and after each injection through 4 cycles, with each cycle comprised of 3 minutes mixing, 2 minutes waiting and 2 minutes measuring. Data was normalized with non-mitochondrial respiration. ATP-linked OCR value was calculated by subtracting basal OCR from proton leak OCR.

### Computational analysis of the single-cell RNA-seq data

For the analysis, an indexed mm10 reference genome was build based on the GRCm38 assembly and genome annotation release 94 from Ensembl. For the alignment, QC, the estimation of valid barcodes and creating the count matrices, the Cell Ranger pipeline (version 2.1.1, from 10xgenomics, https://support.10xgenomics.com) was run with the command “cellranger count” with standard parameters, except that the number of expected cells was set to 10,000, the chemistry was set to “SC3Pv2”. An anndata object was created using the python package Scanpy^33^ (version 1.0.4). The full analysis is shown in **Supplementary Data 1**.

#### Cell type assignment

The workflow was performed in scanpy version 1.4.3 (*55*). Cells were filtered based on a UMI counts and the fraction of mitochondrial RNA. The remaining cell vectors were normalized to sum a total count of 1e4 by linear scaling, log(x+1) transformed and highly variable genes were selected. We then performed PCA with 50 PCs and used the PC space to compute a k-nearest neighbor (kNN) graph (k=100, method=umap). We computed UMAP and a louvain clustering (resolution=1, flavor=vtraag) based on the kNN graph (louvain_1). We assigned cell types to clusters based on marker gene expression by cluster.

#### Heterogeneity analysis of preadipocytes

We selected the louvain clusters that correspond to preadipocytes and putative mature adipocytes from the overall clustering (louvain_1) and separately processed these cells: normalizing to 1e4 counts, then log(x+1) transforming the data and selected highly variable genes followed by PCA with 50 PCs. Again, we computed a kNN graph (k=100, method=umap) based on the PC space. We computed UMAP and a louvain clustering (resolution=1, flavor=vtraag) based on the kNN graph, we call this clustering louvain_2 in the following

### Statistical analysis

Data are shown as mean ± SEM, if not indicated otherwise in the figure legend. Statistical significance for multiple comparisons was determined by One-or Two-Way ANOVA, with Tukey’s multiple comparisons test, or unpaired t-test (two-tailed P value). Correlation graphs were analysed with linear regression (two-tailed P value, 95% CI). GraphPad Prism 6 was used for statistical analysis. P values <0.05 were considered as statistically significant.

RNA count files were normalized using R package DESeq2. Genes where all expression values were in the lowest 25% of the data were removed. Additionally, genes with a variance in the lowest 25% were removed from the data. Hierarchical clustering was performed using ‘Euclidean’ distance measure and nearest distance linkage method, if not indicated otherwise.

Nonlinear dimensionality reduction was performed using Laplacian eigenmaps (*56*) with Mahalanobis distance as similarity measure and *k* = 23 for nearest neighbor graph generation. First eight eigenvectors of the graph Laplacian matrix were used as a low dimensional representation of preadipocyte expression matrix. For each gene in eigenvector-space we calculated the standardized Euclidean distance to geometrical median. K means with *k* = 2 was used to remove ‘uninformative’ proximal genes from the data. Silhouette was used to estimate an optimal *k =* 7 for k-means clustering of the remaining distal genes. KEGG pathway enrichments were calculated using hypergeometrical distribution tests. Network plots were created using Cytoscape v3.6. Calculations were done using R 3.4 and MATLAB 2017b.

## Acknowledgements

This work was supported by the project Aging and Metabolic Programming (AMPro) and by the European Research Council ERC STG (AstroNeuroCrosstalk no. 757393) to CGC.

## Author contribution

R.K., I.A., T.S., A.I, I.K.F., S.K. and A.B. performed experiments. D.L. Analyzed data. C.C., J.W.J and A.M.C. designed, conducted, and analyzed the human tissue. K.S. performed automated imaging, A.F. and A.W. performed quantitative image analyses. D.F., T.W., V.M., M.H., H.L., C.G.C and F.J.T. contributed to the scRNAseq analysis. E.G. and T.S. performed the RNAseq. S.U., D.L. and R.K. designed experiments and wrote the manuscript.

## Data and materials availability

RNAseq data are available at Gene Expression Omnibus (GEO): Series GSE122780 scRNAseq data will be provided to the reviewers upon request and deposited to public repositories upon acceptance.

## Figure legends

**Supplementary Figure 1:**
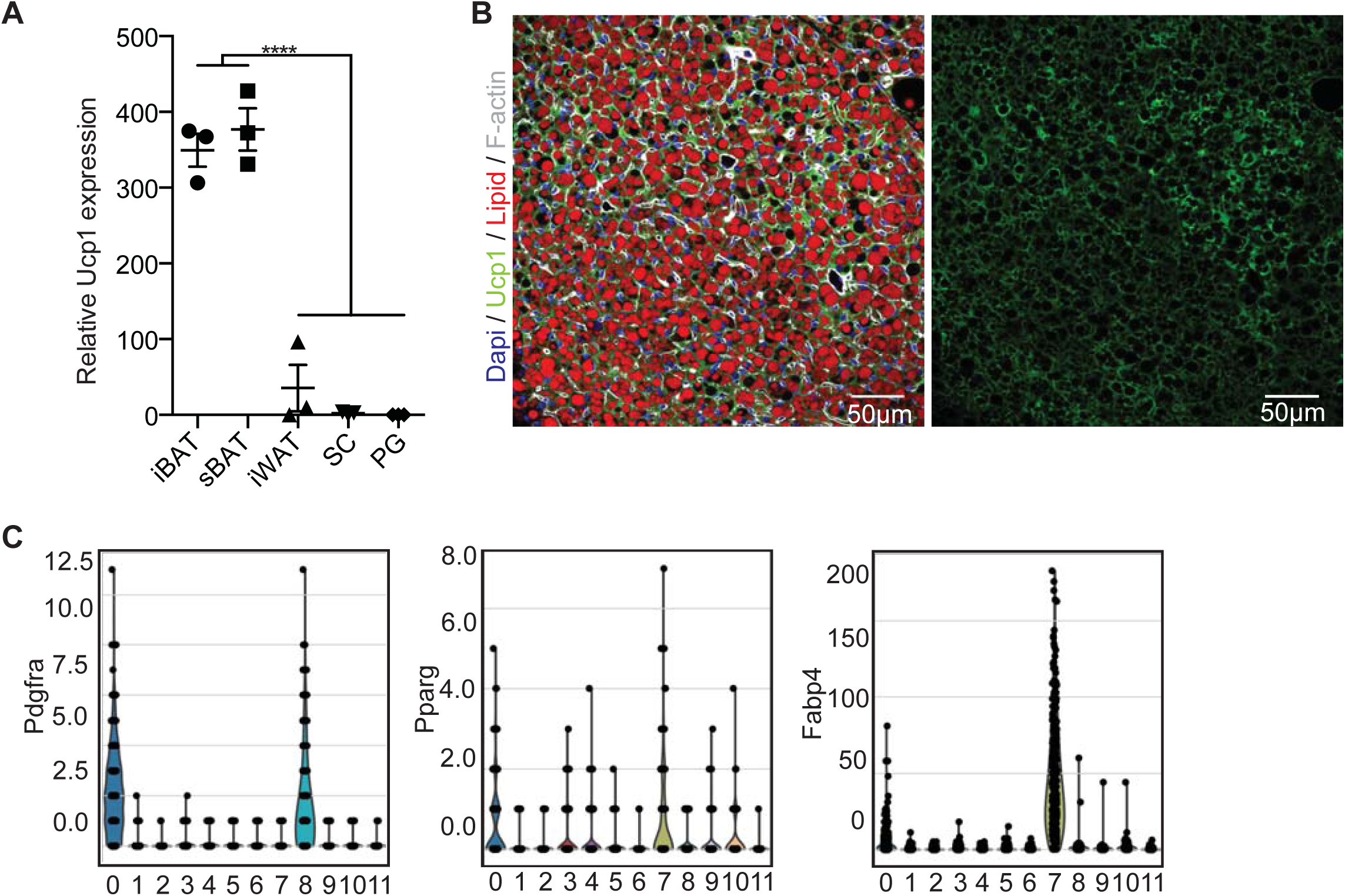
(**A**) Quantitative PCR analysis of Ucp1 expression in interscapular BAT (iBAT), subscapular BAT (sBAT), interscapular white adipose tissue (iWAT), subcutaneous fat (SC) and perigonadal fat (PG) from 6 weeks old C57BL/6 mice (n= 3; *** p< 0.001). Data are mean expressions normalized to TBP ± SEM. (**B**) Representative immunofluorescence staining of Ucp1 on BAT used for quantifications in Figure 1b (DAPI: blue, Ucp1: green, lipid droplets: red, F-actin: gray). (**C**) Violin plots showing the distribution of selected marker genes (Pdgfra, Pparg and Fabp4) across louvain cluster computed on all cell types identified in the single-cell RNA-seq data set.

**Supplementary Figure 2:**
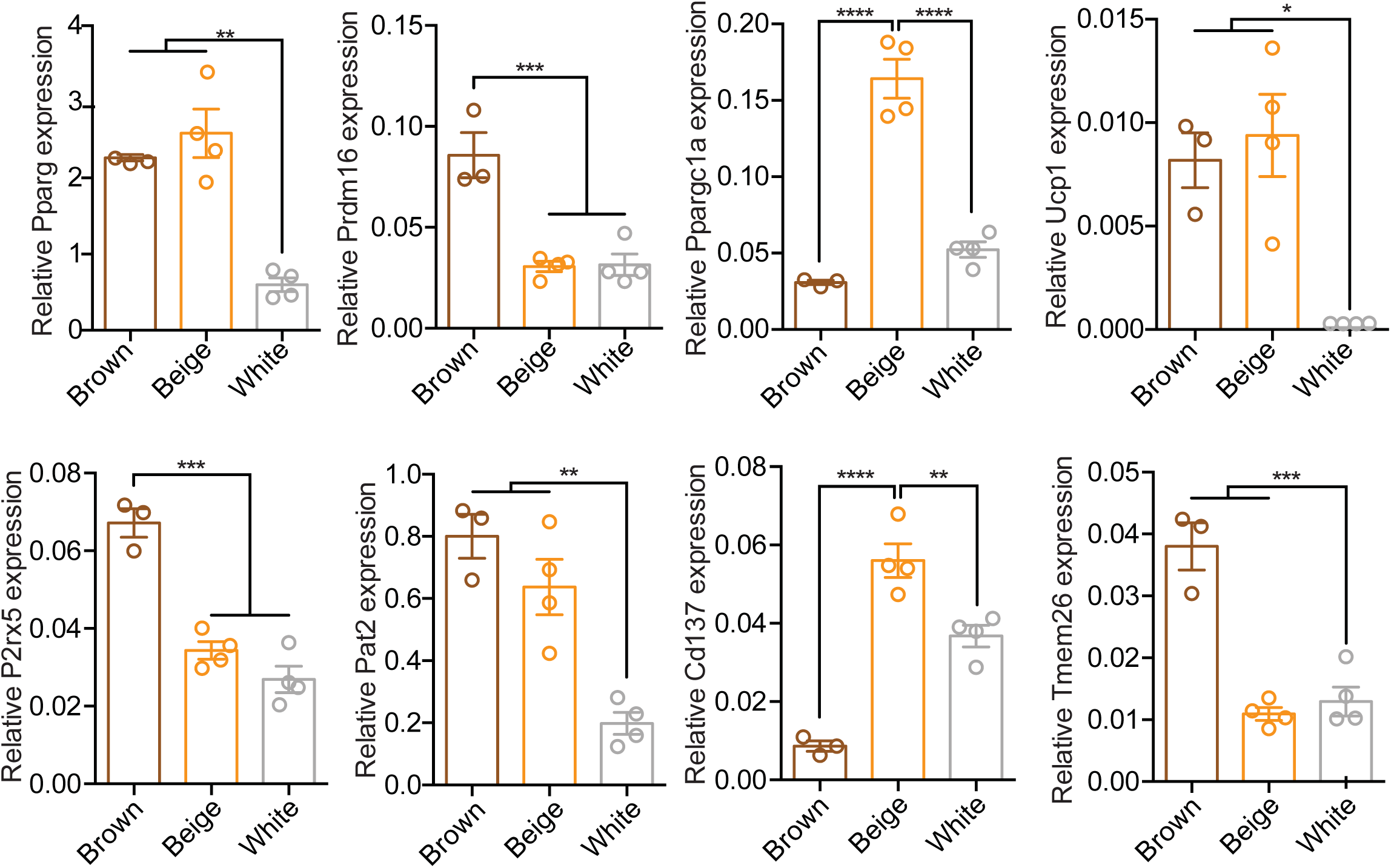
All qPCR data were normalized to TBP and shown as mean ± SEM, (n= 3), * p< 0.05, **, ^##^ p< 0.01, *** p< 0.001, ****p< 0.0001.

**Supplementary Figure 3:**
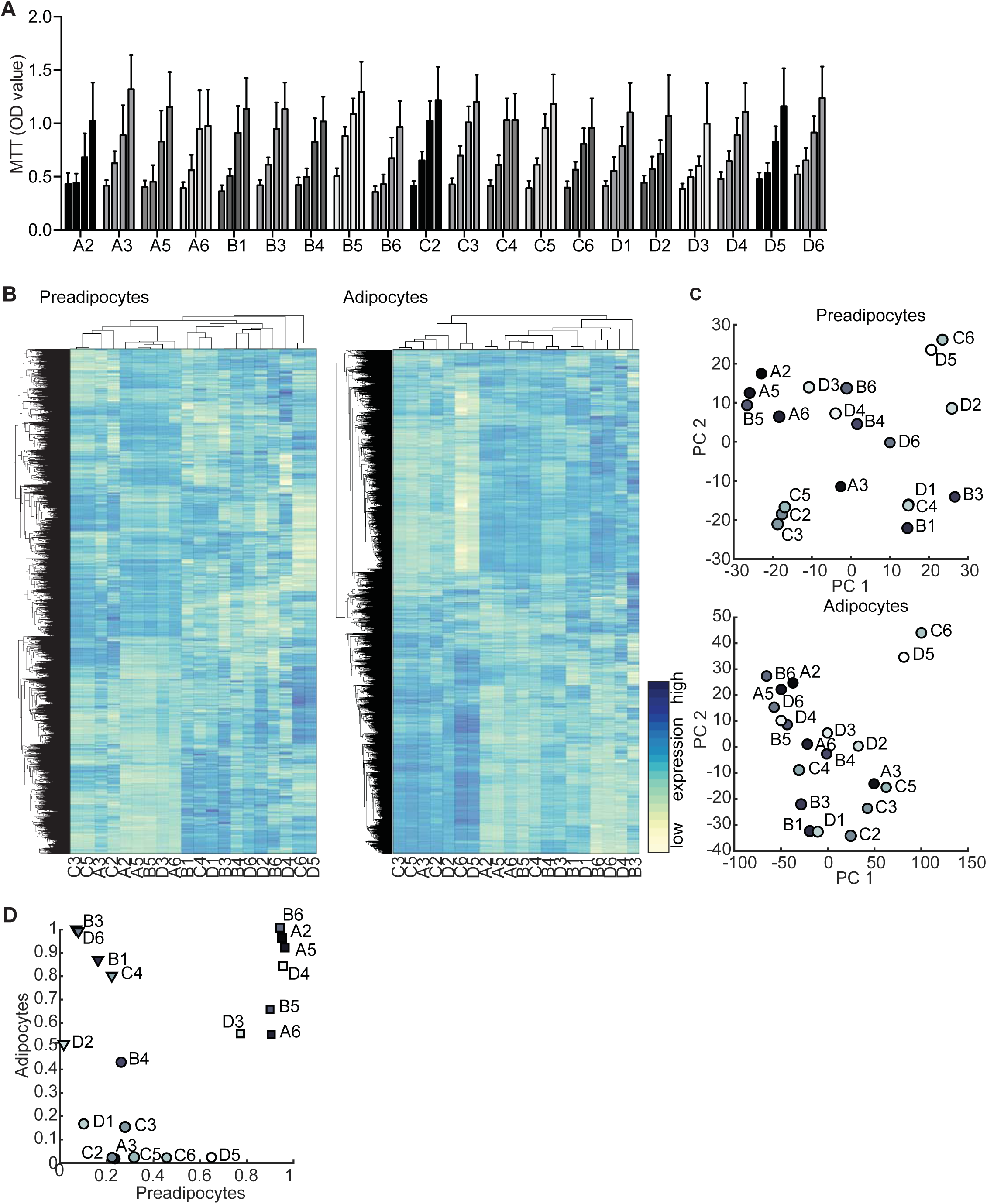
(**A**) MTT assay of 20 preadipocyte clones for 4 days (left to right bar, n=3-5). (**B**) Heatmap of all 9483 expressed genes in preadipocyte. Gene expression in rows was z-score normalized, columns refer to cell lines (left panel). Hierarchical clustering heatmap of all 10363 expressed genes in differentiated adipocytes (right panel). (**C**) PCA plot of the first two PC of pre-adipocytes expression data (upper panel) and differentiated adipocytes (lower panel). (**D**) Scatterplot comparison of the estimated BATness from pre- and differentiated adipocyte cell lines. Dots, triangles and square indicate the three clusters identified by k-means clustering.

**Supplementary Figure 4:**
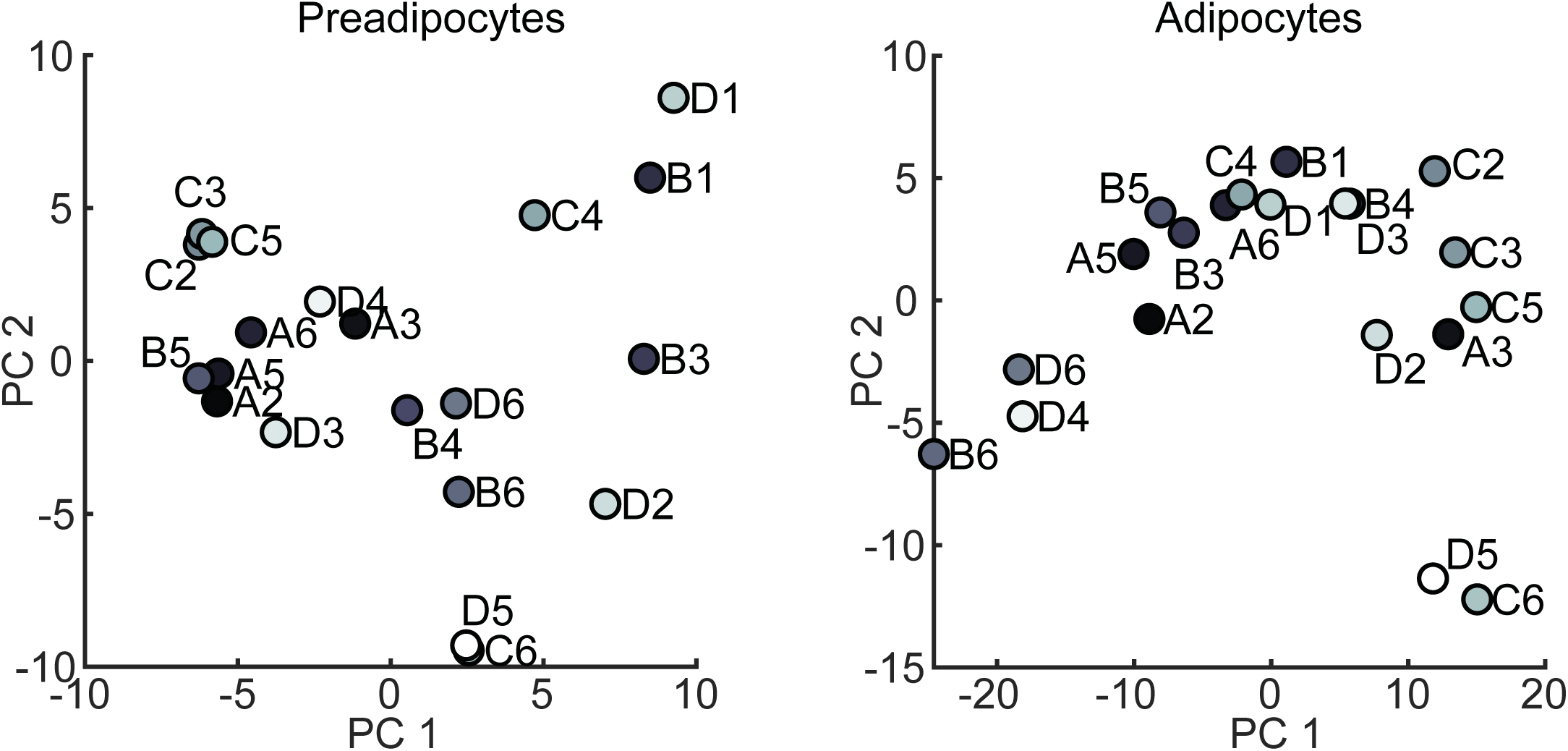
PCA plot for the first two PCs of GEM genes from pre-adipocytes (left panel) and differentiated brown adipocytes (right panel).

**Supplementary Figure 5:**
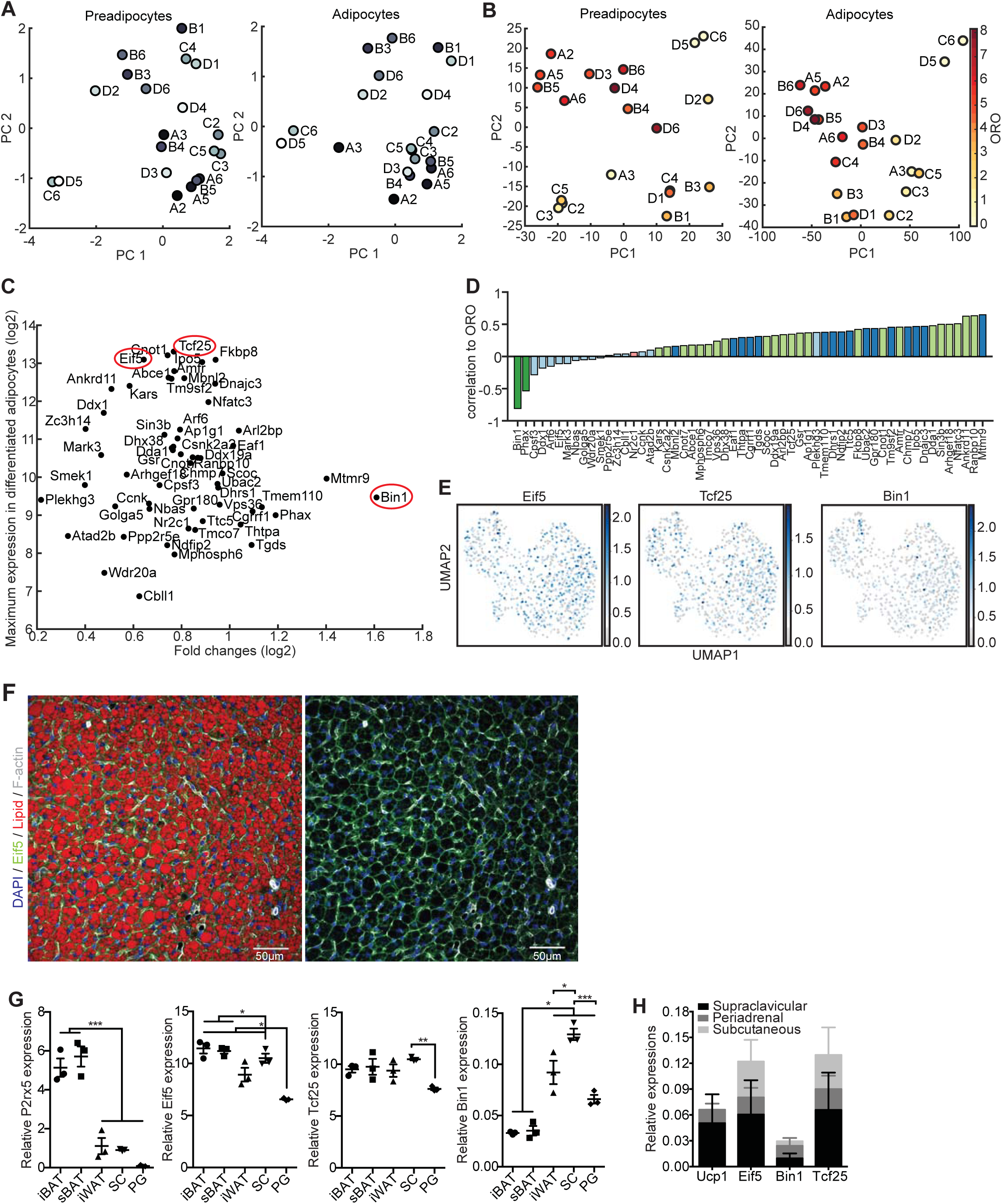
PCA plots of pre-(left panel) and differentiated (right panel) adipocytes (**A**) from stable expressed GEM genes and (**B**) color-coded clones based on the ORO quantification. (**C**) Scatter plots of stable expressed genes with regards to their maximum expression in differentiated adipocytes and maximal log fold change (log2) between differentiated adipocyte clones. (**D**) Correlation coefficients for stably expressed genes compared to lipid content as measured by ORO. Color of bars indicates module membership. (**E**) UMAP of all preadipocytes with Eif5, Tcf25 and Bin1 expression superimposed. The expression values shown are size factor normalized and log transformed. (**F**) Representative immunofluorescence staining of Eif5 on BAT sections (DAPI: blue, Eif5: green, lipid droplets: red, F-actin: gray). mRNA expression of P2rx5, Eif5, Tcf25 and Bin1 in (**G**) iBAT, sBAT, iWAT, SC, and PG of wild-type C57BL/6 mice as assessed by qPCR (n= 3, data are mean expression normalized to B2m ± SEM) * p< 0.05, ** p< 0.01, *** p< 0.001 and (**H**) in supraclavicular, periadrenal and subcutaneous adipose tissue biopsies from 6 human donors as assessed by RNAseq.

**Supplementary Figure 6:**
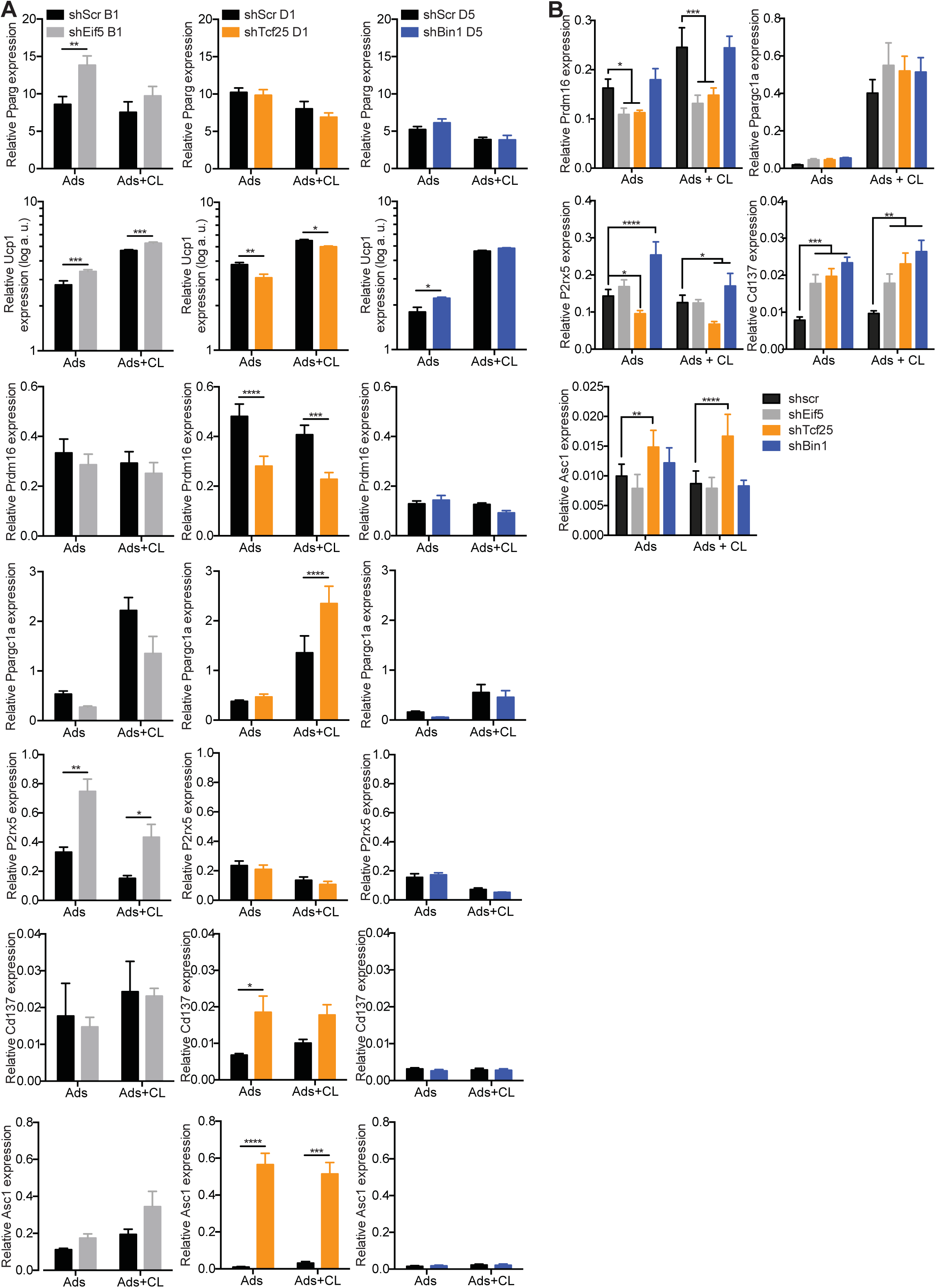
Semi-Quantitative PCR analysis of Pparg, Ucp1, Prdm16, Ppargc1a, P2rx5, Cd137 and Asc1 expression at day 8 of differentiation and 3 hours treatment with 0.5μM CL-316,243 in (**A**) shScr B1 and shEif5 B1 (left panel), shScr D1 and shTcf25 D1 (middle panel), shScr D5 and shBin1 D5 (right panel) cells (n= 5), and (**B**) shScr, shEif5, shTcf25 and shBin1 on mixed brown adipocytes cells (n= 6). Data are shown as mean expression normalized to Tbp ± SEM. * p< 0.05, ** p< 0.01, *** p< 0.001, **** p< 0.0001.

